# Maternal obesity programs white and brown adipose tissue transcriptome and lipidome in offspring in a sex-dependent manner

**DOI:** 10.1101/2021.02.08.430188

**Authors:** Christina Savva, Luisa A. Helguero, Marcela González-Granillo, Tânia Melo, Daniela Couto, Byambajav Buyandelger, Sonja Gustafsson, Jianping Liu, Maria Rosário Domingues, Xidan Li, Marion Korach-André

**Author notes:** Corresponding author: Marion Korach-André, Department of Medicine, Metabolism Unit, Karolinska Institute, S-141 57 Huddinge, Sweden.

## Abstract

The prevalence of overweight and obesity among children has drastically increased during the last decades and maternal obesity has been demonstrated as one of the ultimate factors. Nutrition-stimulated transgenerational epigenetic regulation of key metabolic genes is fundamental to the developmental origins of the metabolic syndrome. Fetal nutrition may differently influence female and male offspring. In this work, we investigated the sex-dependent programming of maternal obesity in visceral, subcutaneous and brown adipose tissues of offspring using magnetic resonance imaging and spectroscopy and a lipidomic approach combined with a Smart-Seq2 differential sequencing analysis. We show that the triglyceride profile varies between adipose depots, sexes and maternal diet. Our results demonstrate for the first time that a sex-dependent gene programming exists in visceral, subcutaneous and brown adipose tissues. Maternal obesity differentially programs gene expression in adipose depots of female and male offspring, which may contribute to the sex-dependent metabolic complications later in life.

**Graphical abstract:** 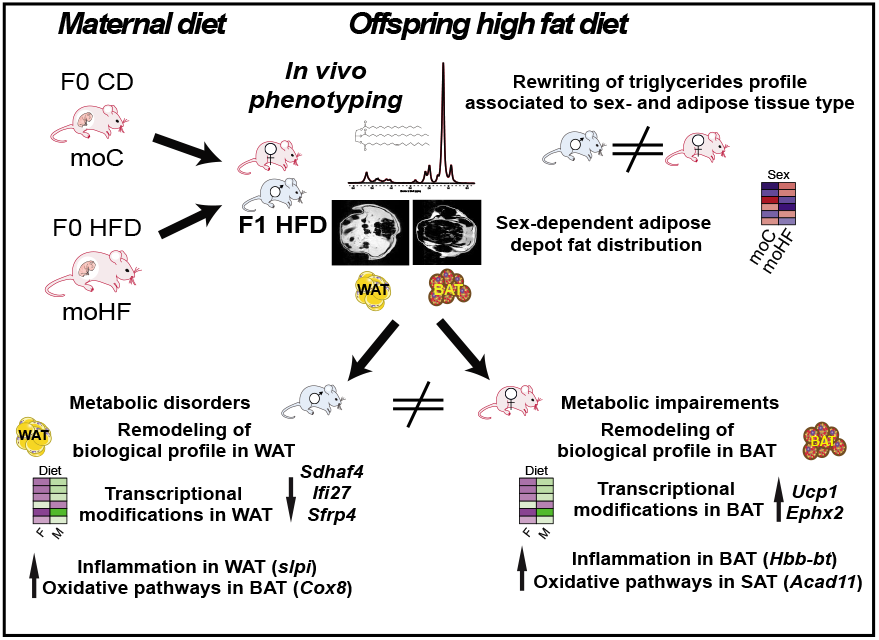

## INTRODUCTION

The drastic increase in consumption of high caloric diets with high levels of modified fat by the food industry, including saturated- and trans-fatty acids associated with a sedentary lifestyle, has dramatically challenged humans’ metabolism worldwide. Metabolic adaptation to these recent lifestyle changes has confronted the scientific community to decipher the impact of nutrients on metabolic homeostasis. Most importantly, the increased prevalence of overweight and obesity in women in reproductive age has urged the need to better understand the impact on the fetus health later in life (Chen et al., 2018). Recently, a large number of studies have demonstrated the noteworthy sensitivity of the offspring to nutritional, environmental and hormonal changes during the prenatal, neonatal and postnatal periods, which facilitates the development of metabolic complications in adulthood (Baker et al., 2017; de Almeida Faria et al., 2017; Frihauf et al., 2016; Gambineri et al., 2020; Krasnow et al., 2011). The intrauterine programming of obesity and associated metabolic risks in offspring adulthood rely on epigenetic regulation as one of the key underlying mechanisms (Elshenawy and Simmons, 2016). Epigenetic modifications induced by maternal obesity (MO) (Deodati et al., 2019; Liang et al., 2016) lead to a cyclical transgenerational transmission of obesity that, in the near future, may become a heavy burden worldwide (Iozzo et al., 2014). Therefore, understanding the link between MO and offspring health would allow to anticipate better public health policy and implement more effective interventions.

Adipose tissue (AT) is a complex and a highly metabolically active organ essential for the regulation of energy balance in order to maintain metabolic homeostasis in the body (Choe et al., 2016). In mammalians, white and brown adipocytes tune energy balance according to the calorie intake and the energy expended. The development of adipose tissue occurs at an early stage, during prenatal and postnatal periods (Desai and Ross, 2011), hence MO may have an impact on programing offspring’s adipose tissue function. Obesity during fetal programming of adipocytes leads to hypothalamic leptin resistance (Dias-Rocha et al., 2018), endocannabinoid signaling dysfunction in white adipocytes (de Almeida et al., 2020) and uncoupling protein-1 dysfunction in brown adipocytes (Dias-Rocha et al., 2018) and hereby promotes development of metabolic diseases later in life in offspring. MO may affect processes in adipose tissue development that can result in adipose tissue dysfunction and lead to adverse effects, promoting metabolic complications.

In the current study, we explored how MO prior to pregnancy, and maintained throughout pregnancy and lactation, can predispose white and brown adipose tissue in offspring fed the high-fat diet (i.e. obese offspring) to metabolic dysfunctions later in life. This was done by characterizing the transcriptome and lipidome of three fat depots: visceral (VAT), subcutaneous (SAT) and brown (BAT) adipose tissue at different time-points in offspring life. Our study also explored the sex-differences in the metabolic response to MO in offspring that may compromise female and male offspring metabolism homeostasis later in life. Our results showed that MO does not affect global adiposity in offspring. However, we observed sex- and adipose tissue-dependent gene regulation in MO offspring that may balance adipose tissues lipidome and physiology and may contribute to the sexual dimorphism observed in the metabolic adaptations in obesity later in life.

## RESULTS

### Maternal obesity alters the physiological and biological adaptations to obesity in a sex-dependent manner

The F0 dam were fed with the high-fat diet (HFD) or the control diet (CD) for 6 weeks prior mating and during pregnancy and lactation. F0 sires were fed with the CD throughout the study. All F1 offspring were fed with the HFD after weaning. HFD-dam (moHF) weighted significantly more than CD-dam (moC) prior mating (Fig.1A). Body weight of offspring was sex dependent but not maternal diet dependent. Males weighted significantly more than females regardless of the maternal diet at midterm (MID, S; p<0.001) and endterm (END, S; p<0.05) (Fig.1B and Table 1). Average food intake was similar between sexes irrespective of the maternal diet, though maternal obesity (MO) tended to induce food intake in female and to reduce it in male offspring (Fig.1C, S; p<0.001 and I; p<0.05). In order to define if MO altered total adiposity in offspring in the short or/and long term, we performed *in vivo* magnetic resonance imaging (MRI) at 15 weeks (MID) and 25 weeks (END) of age using the mouse as its own control (Fig.1D). At MID, males born from obese mothers (M-moHF) accumulated less fat than males born from control mothers (M-moC) (Fig.1E, D; p<0.01, S; p<0.05 and I; p<0.05), which was normalized at END; with females getting more total fat on BW ratio (TF:BW) compared to males regardless of the mother diet (Fig.1E, S; p<0.01). The proportion of fat stored in the visceral region (VAT:TF) was lower than in the subcutaneous region (SAT:TF) in both sexes. At MID, females had lower VAT:TF and higher SAT:TF than males irrespective of the mother diet, however these differences disappeared at END (Fig. 1F-1G). Collectively, our results reveal that MO does not affect the overall fat distribution neither in female nor in male offspring on the long term (END). Interestingly, on a short term (MID) our results indicate that MO diminished total adiposity in males, with males having proportionally more VAT and less SAT than females, which is correlated with metabolic dysfunctions.

**Figure 1.**
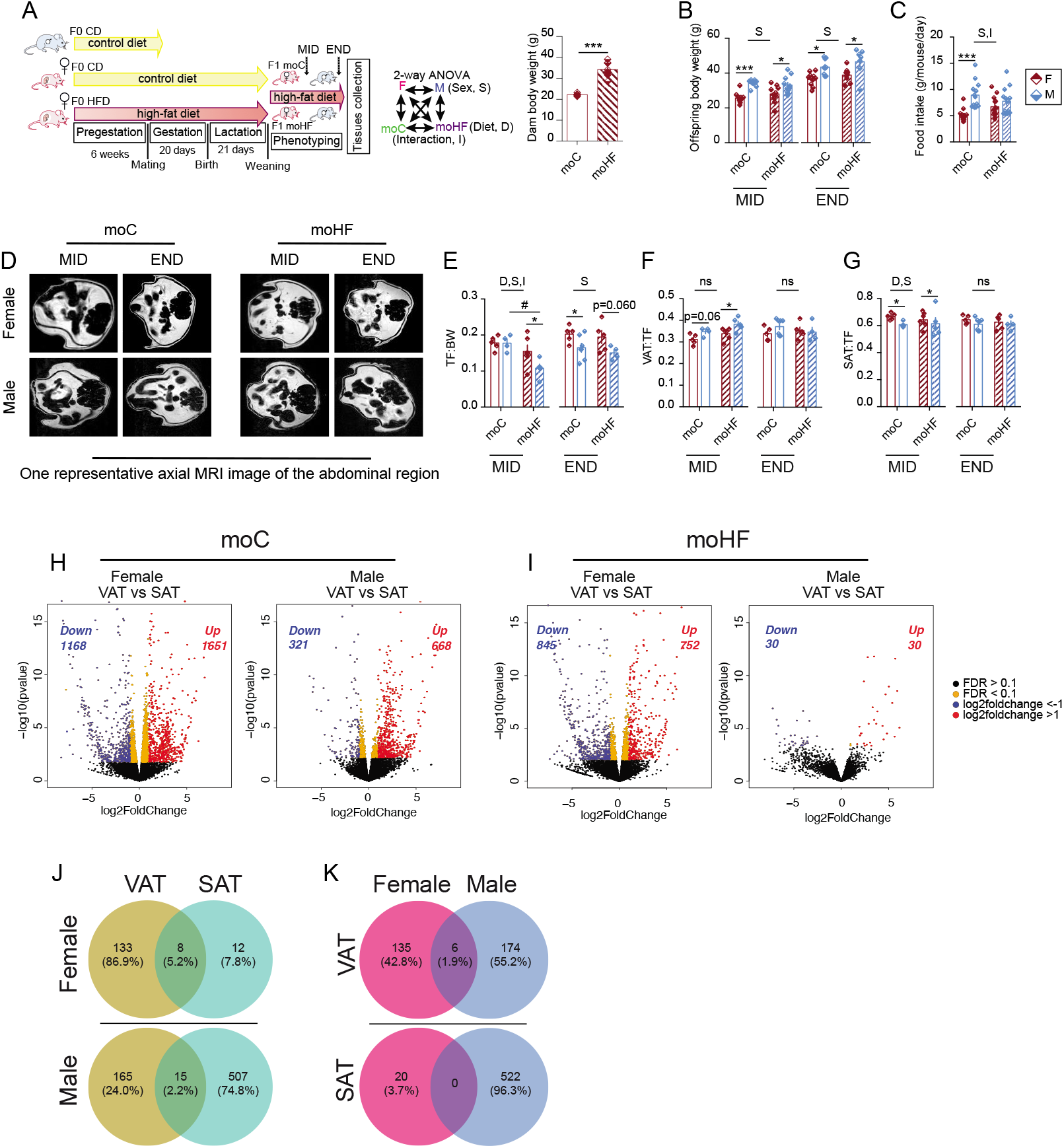
Maternal obesity affects adiposity differently in obesogenic male and female offspring. (A) Experimental setting of the study, two-way ANOVA statistical comparisons and pre-gestational body weight of dam-F0 in CD group (n=6) (open bar) and HFD group (n=6) (stripped bar). (B) Time course body weight curve in female and male offspring during the 27 weeks following birth (n=10-12/sex/diet). (C) Average food intake in offspring. (D) MRI images of the lower abdominal region in female and male offspring. (E) Total fat on body weight (TF:BW) (n=5-7). (F) Visceral fat on TF ratio (VAT:TF) (n=5-7). (G) Subcutaneous fat on TF ratio (SAT:TF) (n=5-7). (H) Volcano plots of Smart-Seq2 data comparing VAT and SAT in moC (females, n=5 and males, n=5). (I) Volcano plots of Smart-Seq2 data comparing VAT and SAT in moHF (females, n=6 and males, n=3). Significantly (FDR< 0.1) upregulated (red dots) and downregulated (blue dots) genes over log2foldchange >1 and <-1. Orange dots indicate the genes that are significantly changed (FDR < 0.1). Black dots indicate genes that are not significant. (J) Venn diagram of all significantly DEG in response to MO in female and male offspring, between VAT and SAT. (K) Venn diagram of all significantly DEG in response to MO in VAT and SAT, between female and male offspring. Two-way ANOVA (sex (S), mother diet (D), interaction (I) between sex and diet, and (ns) for not significant) followed by a Tukey’s multiple comparisons test when significant (P<0.05) in figures B, C, E, F and G. ^*^, males *versus* females and ^#^, moHF *versus* moC (P<0.05); ^**^, P<0.01; ^***^, P<0.001.

**Table 1.**
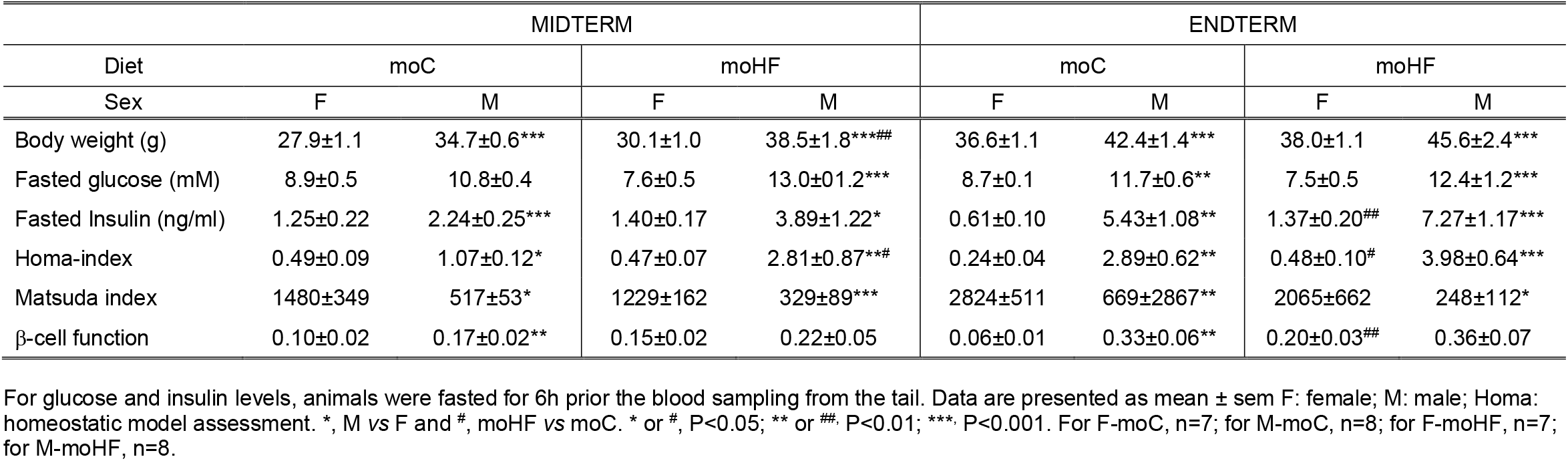
Plasma levels and markers of insulin sensitivity in F and M offspring born from CD (moC) or HFD (moHF) mothers at MID and END term. For glucose and insulin, animals were fasted for 6h prior the blood sampling from the tail. For body weight, F-moC (n=11) and M-moC (n=11), for F-moHF (n=11) and M-moHF (n=10); for the rest F-moC (n=7) and M-moC (n=7), for F-moHF (n=8) and M-moHF (n=7). Unpaired two-tailed Student’s *t*-test were considered significant when P<0.05. ^*^, M *vs* F and ^#^, moHF *vs* moC. ^*^ or ^#^, P<0.05; ^**^ or ^##^, P<0.01; ^***^, P<0.001. F: females; M: males; HOMA: homeostatic model assessment. Body weight and plasma parameters in female (F) and male (M) offspring.

To elucidate the underlying mechanisms by which MO may alter offspring’s adipose metabolism, we performed a Smart-Seq2 differential gene expression analysis in VAT and SAT of female and male offspring in moC and moHF groups (Fig.1H–1I). In females born from control mothers (F-moC), 2,819 (up- and downregulated) differently expressed genes (DEG) were found in VAT *versus* SAT, while in M-moC, only 989 genes were found significantly regulated (Fig.1H). Surprisingly, with MO 1,597 DEG were identified between VAT and SAT in females and only 60 DEG in males (Fig.1I). These results indicate that VAT and SAT have a different gene expression profile, especially in females. In addition, MO drastically remodels gene expression between VAT and SAT in males. To further dissect the impact of MO on the gene expression pattern in offspring’s white adipose depots, we presented all the DEG in Venn diagrams. In females, we identified 133 and 12 DEG exclusively in VAT and SAT, respectively, and 8 DEG common between VAT and SAT, in response to MO (Fig.2J). In males, 165 and 507 DEG were exclusively identified in VAT and SAT respectively, and 15 DEG were common between VAT and SAT, in response to MO (Fig.1J). In conclusion, ~87 % of the DEG by MO in females are changed in the VAT as opposed to males that account for about 75 % of the DEG by MO in the SAT. When we compared the effect of MO on the gene expression profile between sexes in the VAT and in the SAT, we found 135 and 174 DEG in VAT exclusively in females and males, respectively, while only 6 genes were common between sexes (Fig.1K). In SAT, 20 and 522 DEG were exclusively expressed in females and males respectively, whereas no common genes were found between females and males (Fig.1K). To sum, males show stronger transcriptional remodeling by MO especially in SAT compared to females. Surprisingly, very few DEG are shared in response to MO in VAT and SAT between sexes. These results demonstrate that MO affect the transcriptome of female and male offspring in a sex- and tissue-specific manner.

**Figure 2.**
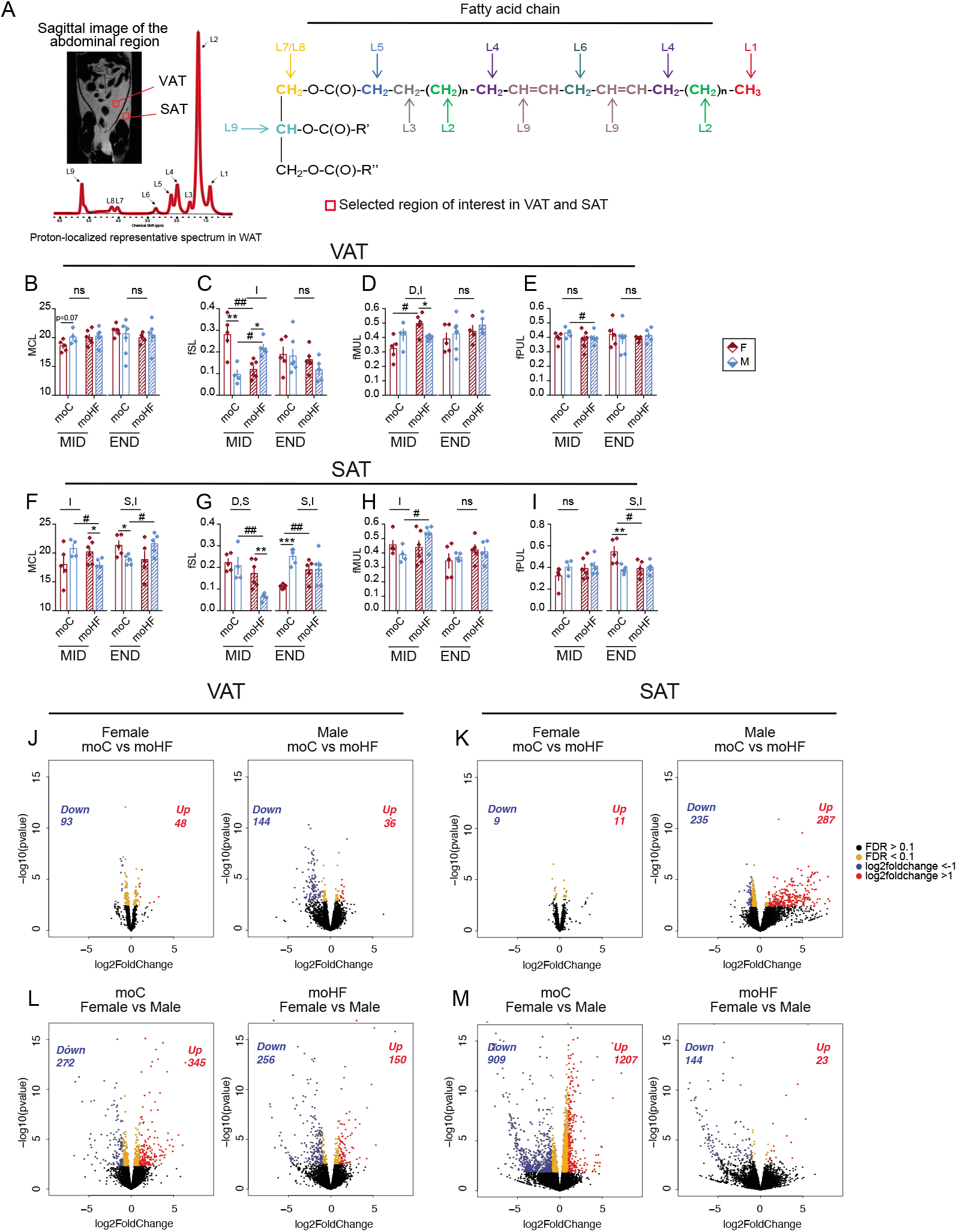
Maternal obesity adjusts triglycerides and gene expression in visceral and subcutaneous adipose tissue in a sex-dependent manner in offspring. (A) Sagittal image of the whole-body fat and one representative ^1^H-localized spectrum for *in vivo* quantification of fatty acids composition of the triglyceride molecule in offspring’s WAT. (B) Mean chain length (MCL) in VAT (n=5-7). (C) Fraction of saturated lipids (fSL) in VAT (n=5-7). (D) Fraction of monounsaturated lipids (fMUL) in VAT (n=5-7). (E) Fraction of polyunsaturated lipids (fPUL) in VAT (n=5-7). (F) Mean chain length (MCL) in SAT (n=5-6). (G) Fraction of saturated lipids (fSL) in SAT (n=5-6). (H) Fraction of monounsaturated lipids (fMUL) in SAT (n=5-6). (I) Fraction of polyunsaturated lipids (fPUL) in SAT (n=5-6). (J) Volcano plots of Smart-Seq2 data comparing moC and moHF in females (n=6) and males (n=3) VAT. (K) Volcano plots of Smart-Seq2 data comparing moC and moHF in females (n=6) and males (n=3) SAT. (L) Volcano plots of Smart-Seq2 data comparing females (n=6) and males (n=3) VAT in moC and moHF groups. Significantly (FDR< 0.1) upregulated (red dots) and downregulated (blue dots) genes over log2foldchange >1 and <-1. Orange dots indicate the genes that are significantly changed (FDR < 0.1). Black dots indicate genes that are not significant. Two-way ANOVA (sex (S), mother diet (D), interaction (I) between sex and diet, and (ns) for not significant) followed by a Tukey’s multiple comparisons test when significant (P<0.05). ^*^, males *versus* females and ^#^, moC versus moHF (P<0.05), ^**^ or ^##^, P<0.01, ^***^, P<0.001.

### Maternal obesity modifies triglycerides composition in visceral and subcutaneous adipose tissue in a sex-dependent manner

Fat distribution is sex-dependent but the underlying mechanism by which triglycerides (TG) are deposited in different parts of the body between sexes remains unknown. Moreover, sex-specific lipid composition in fat depot could trigger the sex-dependent differences in body fat distribution; hence a characterization of these fat depots will shed light into the still unknown mechanism by which lipids are stored in different locations in the body between sexes. Therefore, we performed *in vivo* proton magnetic resonance spectroscopy (^1^H-MRS) in VAT and SAT of offspring at two timepoints (MID and END) to evaluate the TG profile in terms of the fatty acid (FA) length and saturation. Nine lipid signals were identified in the murine adipose tissue ^1^H-MRS spectra and concentrations of each lipid class were derived from the area of the resonance peaks of the individual metabolites as shown in Fig.2A.

No differences of medium chain length (MCL) of the TG in VAT were observed between all groups and at both timepoints (Fig.2B). In contrast, MCL in SAT was different between sexes and MO had a significant impact on its distribution. At MID, MCL was higher in M-moC than in F-moC, and inversely lower in males compared to females in moHF group (I, p<0.05). At END, MO decreased and increased MCL in females and males, respectively; as a result, M-moC had lower MCL than F-moC and higher in moHF (Fig.2F; S,I, p<0.01). The fraction of saturated lipids (fSL) in VAT was higher in F-moC than in M-moC at MID, and MO reduced and increased fSL in females and males, respectively, to a lower level in females than in males (Fig.2C; I, p<0.01). At END, no differences were observed in the fSL between all groups. In SAT, it is interesting to note that the fSL was highly sex- and maternal diet-dependent at both MID and END. MO reduced the fSL in males significantly to a lower level than in females at MID (Fig.2G; D,S p<0.05). At END, the fSL was higher in M-moC than in F-moC but similar between sexes in moHF groups (Fig.2G; S,I p<0.01). The fraction of monounsaturated lipid (fMUL) in VAT was increased with MO in females to a higher level than males at MID and was normalized in all groups at END (Fig.2D; D,I p<0.05). In SAT, fMUL was increased in M-moHF compared to M-moC at MID (Fig.2H; I p<0.05), while no differences between all groups were observed at END. The fraction of polyunsaturated lipid (fPUL) in VAT was similar between sexes in all groups at both MID and END and MO reduced the fPUL in males at MID (Fig.2E). In SAT, fPUL was similar in both sexes at MID, while at END, we observed a higher fPUL in F-moC than in M-moC but MO reduced the fPUL in females to the level of males (Fig.2I; S,I p<0.05). In sum, our *in vivo* results reveal for the first time that MO alters the TG profile in VAT and SAT at short term (MID), but mostly in SAT on the long term (END). We show that there is a sex-dependent lipid profile; at MID, sex-dependent profiles were observed in both fat depots, but sex has a long-term effect (END) on the saturation level in SAT only. Interestingly, MO counteracts the sex effect in SAT by promoting the fSL and decreasing the fPUL in females compared to males. These results reveal important information on the role of sex in the TG profile in response to MO.

To assess metabolic parameters indicative for metabolic dysfunctions, we measured fasting glucose and insulin levels (Table 1). At MID, fasting glucose was similar between sexes in moC group but higher in M-moHF than in F-moHF. Insulin level was higher in males than in females irrespective of the maternal diet but was significantly increased by MO at END in females only. Homeostatic model assessment (Homa) index was significantly higher in males than in females, indicative of lower hepatic insulin sensitivity in males and was slightly impaired by MO in females. Matsuda index, a marker of whole-body insulin sensitivity, was lower in males than in females at both timepoint and diet groups. The ratio AUCins:AUCglc, a marker of β-cells function, was impaired in M-moC compared to F-moC, but only females showed reduced β-cells function with MO at END (Table 1). In sum, sex-differences are observed in the serum metabolic profile irrespective of the mother diet; and serum metabolic profile is slightly impaired by MO in females on a long term.

### Maternal obesity modulates the white adipose transcriptome differently between sexes and between adipose depots

To investigate the effect of MO on WAT transcriptome that might account for the sexual dimorphism observed in metabolism, we performed a differential expression analysis in VAT and SAT between moC and moHF groups (Figs.2J–2K) and between female and male groups (Figs.2L–2M). In VAT, we identified 141 DEG by MO in females and 180 DEG in males (Fig.2J). Surprisingly in SAT, we identified only 20 DEG by MO in females and, 522 DEG in males (Fig.2K). These results indicate that MO alters male’s VAT and SAT transcriptome in a much higher extent than in females, especially in the SAT. To further investigate the sex differences at the transcriptional level, we compared the female transcriptome to male transcriptome in moC and moHF groups in VAT and SAT (Figs.2L–2M). In VAT, 617 and 406 DEG and in SAT, 2,116 and 167 DEG between sexes were identified in moC and moHF, respectively. These results support the dogma that females and males have different lipid distribution due to different transcriptional activity in VAT and SAT. Here, we confirm that MO reprograms the transcription of genes in VAT and SAT in a sex- and adipose depot-dependent manner.

To elucidate metabolic plasticity between sexes and in response to MO, we linked the gene regulation with metabolic pathways. The pathway analysis showed that in VAT, 110 pathways were significantly oppositely regulated in males compared to females in moC, and 151 pathways in moHF (SupplFig.S1A, blue and red). MO significantly regulated 120 pathways in females and 162 pathways in males (SupplFig.S1A, green and purple). In SAT, we identified 151 significantly oppositely regulated pathways in males compared to females in moC, and 110 pathways in moHF (SupplFig.S1B, blue and red). MO significantly regulated 93 pathways in females and 147 in males (SupplFig.S1B, green and purple). These results indicate that in offspring, there is a sex-dependent regulation of biological pathways and that MO differently modulates pathways in VAT and SAT between sexes.

### Maternal obesity alters triglycerides composition in the brown adipose tissue of female offspring

Adiposity is balanced depending on energy intake and energy expenditure. Energy intake tended to be modified by MO, but with unchanged global adiposity. WAT has the ability to store large amount of TG contained in a single large cytoplasmic lipid droplet. Brown adipose tissue (BAT) is distinguished from WAT by a high level of mitochondria containing uncoupling protein-1 (UCP1), a unique protein disconnecting the oxidative phosphorylation from ATP synthesis and inducing thermogenesis. The activation of thermogenic adipocytes has a major impact on local and systemic energy balance. Therefore, we investigated the BAT by MRI (Fig.3A). At both MID and END, males showed larger BAT and higher ratio of BAT:TF than females, irrespective of the maternal diet (Figs.3B–3C; S, p<0.001). Irrespective of the maternal diet, males displayed about 20% more TG content in BAT than females, which is associated to decreased insulin sensitivity (Raiko et al., 2015), and in line with the higher expression level of *Cd36*, *Plin2* and *Fabp1* in males compared to females (Fig.3D; Suppl.TableS1). In obesity, modulation of BAT activity and TG profile has been associated to insulin sensitivity (Raiko et al., 2015) and to be sex-specific (Hoene et al., 2014). Therefore, we performed an LC-MS analysis of BAT and we quantified 10 TG classes. TG classes were classified based on abundance into low (TG40, TG42, TG44 and TG58), moderate (TG46, TG48 and TG56) and high (TG50, TG52 and TG54) (Figs.3E–3G). Within the low abundant group, 3 TG classes (TG40, TG42 and TG44) were enriched in F-moC compared to M-moC but these differences disappeared in the moHF group, due to an increase abundance of TG42 and TG44 in males and a decrease abundance of TG40 in females by MO (Fig.3E). Moderate TG46 and TG48 were more abundant and TG56 less abundant in F-moC than M-moC but these differences vanished with MO (Fig.3F). In females, MO reduced TG48 and induced TG56 relative content compared to moC (Fig.3F). Within the high abundant TG classes, higher TG50 and lower TG52 levels in F-moC than M-moC were found, and TG54 was increased in by MO in females with no sex differences in moHF group (Fig.3G). These results show that in addition to sex differences, MO selectively modulates TG classes in BAT especially in females, that tends to counteract the sex-dependent profile determined by CD mothers. We next carefully dissected the 54 TG species that were detected by LC-MS (Fig.3H and Suppl.FigsS2A-2C). We observed 17 of 54 (31 %) significant differences in the relative levels of TG species between sexes in moC group but only 2 of 54 (3 %) in moHF group. These changes observed between the two maternal diet offspring groups were the result of a remodeling of TG species with MO in females (15 of 54) but to a much lesser extent in males (4 of 54); as favored by increased expression level of *Lpl, Pltp* and *Elovl3* genes by MO in females only (Suppl.TableS2). As some metabolic diseases are associated with the degree of hydrogen atom bond saturation within the FA contained into TG molecules, we next inspected the level of TG saturation. We observed that the saturation status was altered by MO in females only. In moC group, the relative level of saturated TG and TG containing MUFA was higher in females than in males and, TG containing 3-double bonds were higher in males than in females (Fig.3I). F-moHF had less TG containing 0-, 1- and 2-double bonds and more of the TG containing 3- and 4-double bonds than F-moC, in line with the increase expression level of *Elovl3* and *Lpl* genes in females by MO (Suppl.TableS2). These results provide unique information on the sex-dependent adaptation to MO in BAT metabolism. These sex differences may contribute to the sexual dimorphism in the metabolic response to MO in obese offspring, in particular in term of insulin sensitivity. FA, as signaling molecules may contribute to metabolic dysfunction in obesity. Among the 10 FA species contained into the TG molecules detected by GC-MS, only C18:3ω6 FA level was different between sexes in moC group and MO remodeled 2 of 10 FA species in both sexes in opposite directions (Suppl.Fig.S3A). Total ω3 FA levels were higher in F-moHF than in M-moHF and ω6 FA level were higher in F-moC than in M-moC. Consequently, the ratio of ω6:ω3 FA was higher in F-moC than in M-moC and inversely, lower in F-moHF than in M-moHF (Suppl.Fig.S3B). No sex differences in the FA saturation profile were found, but MO induced the relative level of PUFA in males (Suppl.Fig.S3C). In sum, females show higher proportion of saturated and MUFA TG and lower PUFA as compared to males. MO targets mostly females to counterbalance this sex differences. The ω6:ω3 ratio is higher in females than males but MO, by increasing omega 3 inverts this ratio.

**Figure 3.**
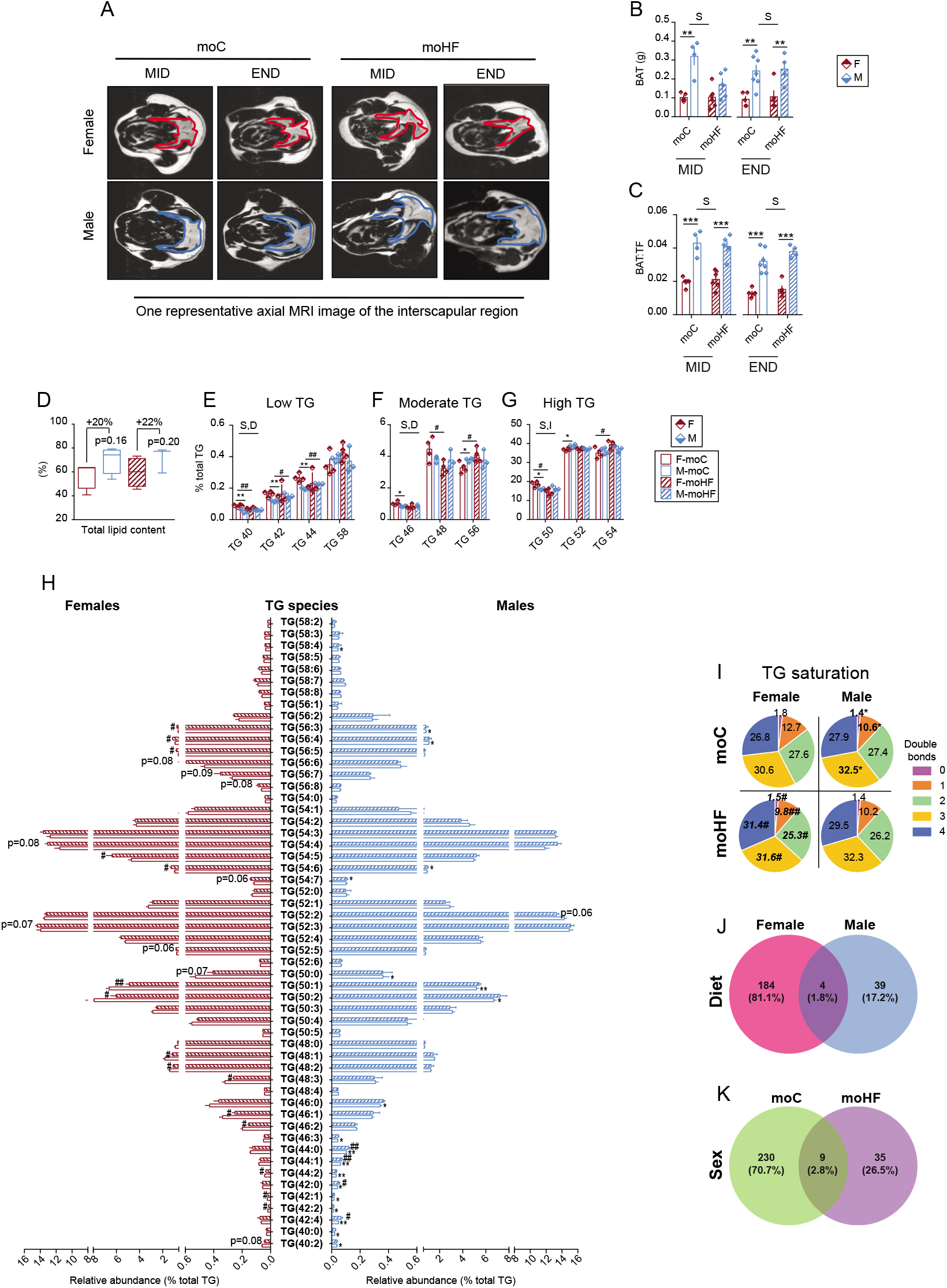
Females brown adipose tissue metabolism is altered by maternal obesity. (A) MRI images of the interscapular brown adipose tissue (BAT) in female and male offspring. (B) BAT quantification based on MRI images (n=5-7). (C) BAT on total fat (BAT:TF) ratio (n=5-7). (D) Fat percentage in BAT (n=4). (E) Low abundant TG classes in BAT (n=4). (F) Moderate abundant TG classes in BAT (n=4). (G) High abundant TG classes in BAT (n=4). (H) TG species detected by LC-MS in BAT. (I) TG saturation profile in BAT. (J) Venn diagram of all significantly DEG between moC and moHF in offspring BAT. (K) Venn diagram of all significantly DEG between female and male offspring in BAT. Two-way ANOVA (sex (S), mother diet (D), interaction (I) between sex and diet, and (ns) for not significant) followed by a Tukey’s multiple comparisons test when significant (P<0.05) in figures B, C, E, F and G. ^*^, males *versus* females and ^#^, moHF *versus* moC (P<0.05); ^**^, P<0.01; ^***^, P<0.001.

### Maternal obesity alters gene expression in the brown adipose tissue of offspring in a sex-dependent manner

To further dissect the sex differences and the impact of MO on offspring’s BAT biology, we performed SmartSeq2 analysis and identified 184 DEG in females and 39 DEG in males in response to MO. Only four genes were shared between sexes in response to MO (Fig.3J). 230 DEG between sexes in moC and 35 in moHF were found, whereas 9 DEG between sexes were common in moC and moHF (Fig.3K). In conclusion, ~80% of the regulated genes by MO in females are changed in the BAT as opposed to males that showed only about 17% of DEG by MO. Sex differences in DEG in BAT account for ~70% in moC but only 26% in moHF. In sum, DEG in BAT is highly sex-dependent in offspring-moC. MO provokes an important remodeling of gene expression in female BAT that may alter metabolism homeostasis as a result. We next inspected the biological pathways that may be sex- or maternal diet-dependent. We found 82 pathways differently regulated between sexes in moC, and 159 in moHF groups (SupplFig.S1C, blue and red). Moreover, MO significantly regulated 126 pathways in females and 138 in males but most of them in opposite directions (SupplFig.S1C, green and purple). These major findings indicate that there is a sex- and maternal diet-dependent regulation of biological pathways in BAT of offspring and that MO modulates oppositely between sexes a large majority of the KEGG pathways in BAT.

### Maternal obesity increases offspring susceptibility to inflammation and insulin resistance in an adipose tissue- and sex-specific manner

To link the physiological and biological adaptations to MO and the sex differences to transcriptional modifications in offspring, we extracted 18 key KEGG pathways that drive metabolism in obesity (i.e. insulin and glucose metabolism, inflammation, oxidative phosphorylation and lipid metabolism) in VAT, SAT and BAT. In VAT, insulin resistance and signaling were upregulated in males and glucose pathway was downregulated in females by MO (Fig.4A), in contrast to SAT where MO downregulated insulin signaling and glucose pathways in males (Fig.4B). In BAT, insulin pathways were highly sex- and maternal diet-dependent (Fig.4C). In both diet conditions, females showed higher expression level of insulin pathways than males, in line with the higher expression levels of *Irs1, Aldoa, Aldob, Hk2* and *Pdha1* in females compared to males (Suppl.Tables S1-S2). MO upregulated insulin secretion but downregulated insulin signaling in females; in contrast MO downregulated all insulin pathways in males. Glucose pathways were upregulated in F-moC compared to M-moC, while MO down- and upregulated the pathways in females and males, respectively (Fig.4C). Obesity is associated with low grade, chronic inflammation that in turn, will promote the development of insulin resistance and diabetes. Inflammatory pathways were upregulated in males VAT compared to females regardless of the mother diet, and MO increased inflammatory pathways expression in males only. In SAT, inflammatory pathways were upregulated in F-moC compared to M-moC, but MO increased the inflammatory pathways only in males to a significantly higher level than F, as supported by the gene expression (Suppl.Tables S1-S2). In BAT, inflammatory pathways were upregulated in M-moC compared to F-moC, and MO oppositely regulated them between sexes leading to higher inflammation in females, in line with the increased expression of *Ccl9, Cxcl12, Tyrobp* and *Cebpb* inflammatory genes (Suppl.Table S2). In VAT, oxidative phosphorylation pathway was reduced by MO in both sexes, with higher level in males. In SAT, oxidative pathways were both higher in males compared to females independently of the mother diet. MO upregulated oxidative phosphorylation pathway but down regulated the citrate cycle (TCA) pathway in females, and remarkable downregulated all oxidative pathways in males, in line with the gene expression pattern (Suppl.Table S1). In BAT, oxidative pathways were oppositely regulated between sexes and in response to MO. There were enhanced and repressed in females compared to males in moC and moHF groups, respectively. Interestingly in VAT, MO upregulated the lipolysis and reduced PPAR signaling and fatty acid degradation pathways in both sexes. In SAT, lipid metabolism pathways were higher expressed in M-moC than in F-moC; MO repressed most of them only in males. In BAT, MO upregulated the fatty acid pathways but downregulated the lipolysis pathway in males. No effect of MO was observed on females.

**Figure 4.**
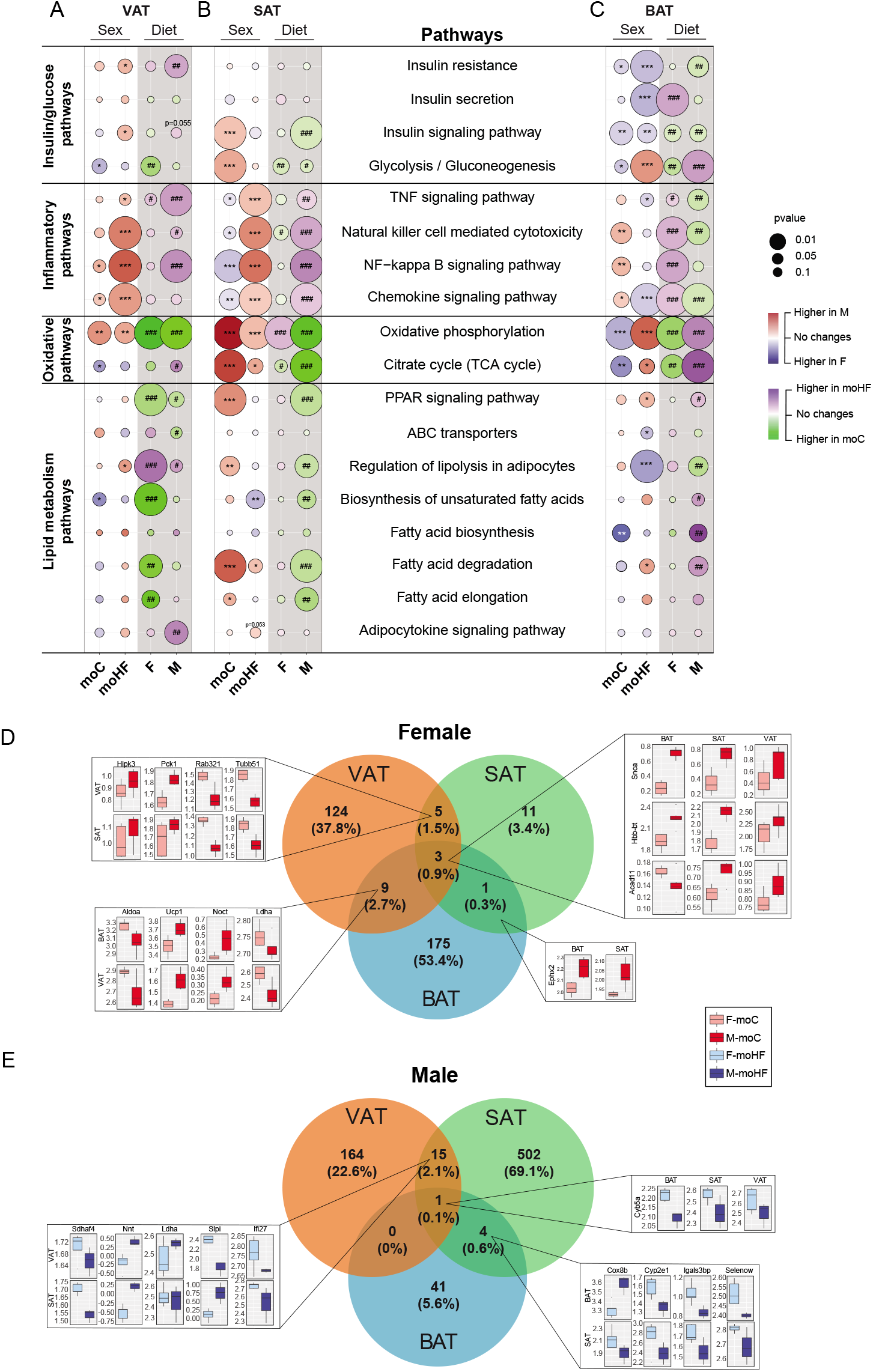
Maternal obesity reprograms metabolic pathways and gene expression in adipose tissue of offspring. (A) Bubble charts showing the sex-dependent and maternal diet-dependent regulation of key KEGG metabolic pathways in VAT (n=3-6). (B) Bubble charts showing the sex-dependent and maternal diet-dependent regulation of key metabolic pathways in SAT (n=3-6). (C) Bubble charts showing the sex-dependent and maternal diet-dependent regulation of key metabolic pathways in BAT (n=3-6). (D) Venn diagram all significantly DEG between moC and moHF in VAT, SAT and BAT of female offspring. Boxplots of the selected significant genes (n=3-6). (E) Venn diagram of all significantly DEG between moC and moHF in VAT, SAT and BAT of male offspring. Boxplots of the selected significant genes (n=3-6). The white background in the bubble charts represents the sex comparisons and the grey background the mother diet comparisons. The color of the bubbles indicates the expression level of the pathways between the groups where red indicates upregulation in males and blue upregulation in females in moC and moHF groups (sex effect). The green color indicates upregulation in moC and purple indicates upregulation in moHF in females and males (maternal diet effect). The size of the bubble indicates the significant level where the bigger the bubble the lower the p-value (higher significance). F: females and M: males.

To further explore which genes might contribute most to the observed changes in the metabolic pathways in VAT, SAT and BAT we obtained the differently regulated genes by MO between the three tissues in females and males. In females, 37.8%, 11% and 53.4% DEG were exclusively found in VAT, SAT and BAT, respectively. Three DEG were common in the three tissues, one DEG was shared between SAT and BAT, while nine DEG were shared between VAT and BAT and five were common between VAT and SAT (Fig.4D). The three genes identified in the three adipose tissues were *Snca* (involved in membrane trafficking), *Hbb-bt* (involved in inflammation) and *Acad11* (involved in FA β-oxidation). Interestingly, *Acad11* was oppositely regulated by MO between BAT (downregulated) and SAT and VAT (upregulated). Among the common genes detected between VAT and SAT we found four genes involved in metabolic pathway, *Hipk3* (involved in steroidogenic pathway), *Pck1* (regulates gluconeogenesis), *Rab32* (involved in mitochondria fusion) and *Tubb51* (involved in obesity). Among the joint regulated genes by MO between VAT and BAT, we extracted *Aldoa* and *Ldha* (which regulate glucose metabolism, downregulated), *Ucp1* (regulates thermogenesis, upregulated) and *Noct* (promotes adipogenesis, upregulated). The single common modulated gene between BAT and SAT was *Ephx2* (involved in ATP metabolism) and was induced by MO in both tissues.

In males, 22.6%, 69.1% and 5.6% DEG by MO were exclusively found in VAT, SAT and BAT, respectively (Fig.4E). The only regulated gene shared by the three adipose tissues, *Cyp5a* (involved in desaturase activity), was downregulated by MO. The four common regulated genes between BAT and SAT, *Cox8b* (regulates mitochondria oxidation), *Cyp2e1* (involved in NAFLD), *Lgals3bp* (regulates inflammation) and *Selenow* (involved in oxidative stress) were all downregulated, except for *Cox8b* that was upregulated only in BAT. 15 regulated genes by MO were found common between VAT and SAT in males, among which, five genes *Sdhaf4* (protects against reactive oxygen species, downregulated) and *Nnt* (involved in mitochondrial electron and proton transports, upregulated), *Ldha* (involved in glycolysis, upregulated), *Slpi* (regulates adipose tissue inflammation, up-in VAT and downregulated in SAT) and *Ifi27* (regulates browning of adipocytes, downregulated) drew our attention due to their key role in metabolism homeostasis and inflammation. These results show that transcriptional regulation by MO is sex- and adipose tissue-specific and may be key contributors to the sex-dependent metabolic adaptation to MO. Interestingly, these results indicate that MO remodeled gene expression mostly in VAT and BAT in females and in VAT and SAT in males.

### Sex-specific genes drive metabolic pathways in VAT, SAT and BAT

To identify the genes involved in the selected signaling pathways in obesity we performed a Chord plot to visualize the biological processes (KEGG) enriched by sexes and by MO. The biological processes included insulin/glucose pathways, inflammatory pathways, oxidative phosphorylation and lipid metabolism along with the log2fold change of each gene in VAT, SAT and BAT (Figs.5A–5C). Evidence of important sexual dimorphism in gene expression in offspring born from lean mothers were observed in all adipose depots. Interestingly, MO suppressed these striking sex differences by remodeling transcriptional activity in the SAT in males and in the BAT in females.

**Figure 5.**
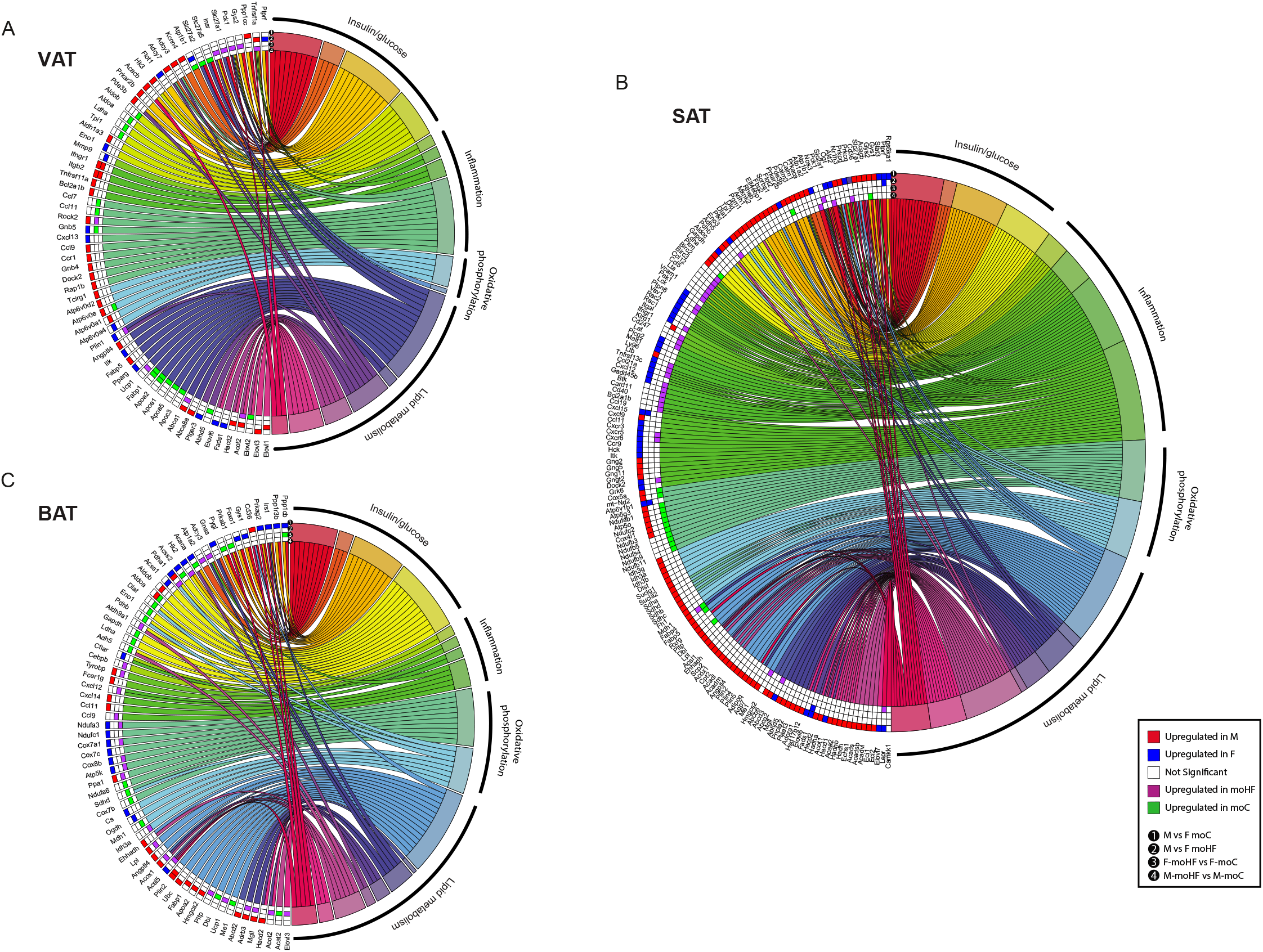
Maternal obesity adjusts gene expression at the transcriptional and post transcriptional levels in VAT, SAT and VAT in a sex- and tissue-dependent manner. (A) Chord plot in VAT demonstrating the DEG clustered into selected pathways and the expression level. (1) female *versus* male (sex) in moC, (2) female *versus* male (sex) in moHF, (3) moC *versus* moHF (diet) in female and (4) moC *versus* moHF (diet) in male (n=3-6). (B) Chord plot in SAT demonstrating the DEG clustered into selected pathways and their expression level. (1) female *versus* male (sex) in moC, (2) female *versus* male (sex) in moHF, (3) moC *versus* moHF (diet) in female and (4) moC *versus* moHF (diet) in male (n=3-6). (C) Chord plot in BAT demonstrating the DEG clustered into selected pathways and their expression level. (1) female *versus* male (sex) in moC, (2) female *versus* male (sex) in moHF, (3) moC *versus* moHF (diet) in female and (4) moC *versus* moHF (diet) in male (n=3-6). Red boxes indicate upregulation in males and blue boxes upregulation in females. The purple boxes indicate upregulation in moHF and green boxes indicate upregulation in moC. White boxes when not significant.

### Maternal obesity drives oppositely the transcriptome between females and males in adipose tissues

The differences in the gene expression profiles in VAT, SAT and BAT of female and male offspring and in response to MO imply that multiple genes are related to several biological pathways. We found that 10 pathways in VAT (Fig.6A), 41 in SAT (Fig.6B) and 23 in BAT (Fig.6C) were significantly and oppositely regulated between females and males in both diet groups (blue and red heatmaps). We extracted 12 pathways in VAT, 32 in SAT and 72 in BAT that were oppositely altered by MO in female and male offspring (green and purple heatmaps). Given that MO altered a large number of pathways between males and females vital for metabolic homeostasis such as fat digestion and absorption, AMPK signaling and inflammatory pathways and steroid hormones biosynthesis, we sought to explore genes that were oppositely expressed between sexes. Surprisingly, we found only two genes (*Dgat2*, involved in TG synthesis and *Sfrp4*, involved in adipogenesis) in VAT (Fig.6D), one gene (*Fabp4*, involved in FA uptake, transport and metabolism) in SAT (Fig.6E) and three genes (*Gbe1*, involved in glycogen degradation, *Idi1*, involved in cholesterol synthesis and *Kng2*, involved in inflammation) in BAT (Fig.6F) oppositely regulated by MO between females and males. These results indicate that a large number of biological pathways appears to be oppositely regulated by MO and between sexes although only few genes are significantly differently regulated. In addition, we show that MO alters differently the white and brown adipose transcriptomes in a sex- and adipose depot-dependent manner. The sex-specific transcriptome in offspring AT may contribute to the sexual dimorphism in obesity and associated metabolic dysfunctions.

**Figure 6.**
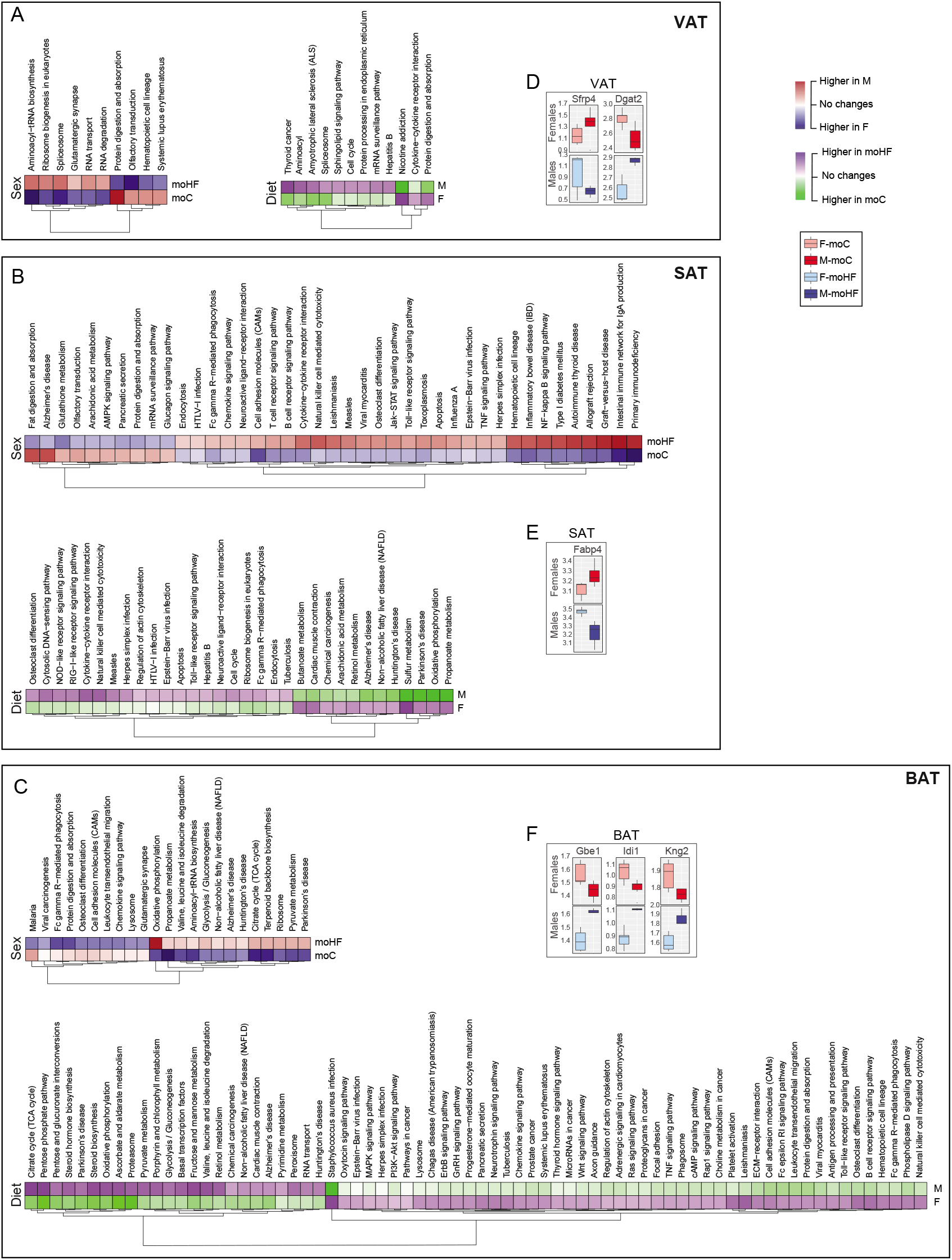
Maternal obesity drives oppositely metabolic pathways in adipose tissues between sexes. (A) Heatmap of significant pathways that were oppositely regulated between female and males (sex) and between moC and moHF (diet) in VAT. Boxplots of the oppositely regulated genes by MO between females and males (n=3-6). (B) Heatmap of significant pathways that were oppositely regulated between female and males (sex) and between moC and moHF (diet) in SAT. Boxplots of the oppositely regulated genes by MO between females and males (n=3-6). (C) Heatmap of selected significant pathways that were oppositely regulated between female and males (sex) and between moC and moHF (diet) in BAT. Boxplots of the oppositely regulated genes by MO between females and males (n=3-6). (D) Selected significantly (FDR < 0.1, P<0.05) oppositely regulated genes by MO between sexes in VAT. (E) Selected significantly (FDR < 0.1, P<0.05) oppositely regulated genes by MO between sexes in SAT. (F) Selected significantly (FDR < 0.1, P<0.05) oppositely regulated genes by MO between sexes in BAT. ^*^, males *versus* females and ^#^, moC *versus* moHF (P<0.05), ^**^ or ^##^, P<0.01, ^***^ or ^###^, P<0.001. F: females and M: males.

## DISCUSSION

In 2016, the World Health Organization estimated that 1.9 billion adults worldwide were overweight and about 650 million were obese and it was anticipated that about 40 −50 % of the women of reproductive age are overweight or obese worldwide. The recent discovery of the changes in the transcriptional and posttranscriptional pathways *in utero* and during lactation (Monks et al., 2018), that increase the susceptibilities to metabolic disorders later in life (Deodati et al., 2019; Seki et al., 2017) is of enormous importance considering the dramatic increased prevalence of obesity in women of reproductive age. In this study, we show that stressing dams with HFD during preconception, gestation and lactation periods has important effects on the developmental programming of white and brown adipose tissues in F1 offspring, in a sex- and adipose depot-dependent manner. These changes may contribute to adipose tissue dysfunction and determine the risk for developing metabolic complications later in life. While several studies have demonstrated the deleterious effect of MO in offspring metabolism (Lecoutre et al., 2016; Liang et al., 2016; Litzenburger et al., 2020; Monks et al., 2018; Sellayah et al., 2019), the novelty of the present study is the combination of advanced physiological *in vivo* and molecular *ex vivo* techniques to further dissect the sex-dependent changes exerted by MO on the offspring adipose metabolism. In addition, we demonstrate complex transcriptional and posttranscriptional mechanisms by which these biological and physiological adaptations are sex- and adipose depot-dependent which may predispose differently females and males to metabolic alteration later in life.

The embryo and fetus development are highly sensitive to its biological environments, and in late gestation, many fetal homeostatic adaptations can be readily perceived. Fetal nutrition (intrauterine environment) and maternal nutrition are not identical. Fetal nutrition lies at the end of a long supply line extending from the maternal macroenvironment through the maternal gastrointestinal, uteroplacental unit and metabolic physiology (Bloomfield and Harding, 1998). Feeding mouse mothers with HFD before mating, during pregnancy and lactation does not lead to significant changes in body weight in female and male offspring compared to offspring born from lean mothers, when offspring fed the HFD postweaning. When the storage capacity of SAT is reduced, in high caloric overload, it leads to fat accumulation in ectopic tissues including VAT and BAT, a phenomenon that is defined as “lipotoxicity” and promotes insulin resistance (Ros Perez and Medina-Gomez, 2011) and low-grade inflammation (Suganami et al., 2012). In the current study, males showed lower SAT and higher VAT accumulation, as well as ~20% higher fat accumulation in BAT than females in both mother diet groups. We showed that females redistribute TG profile in BAT while males redistribute TG profile mainly in SAT in response to MO. These major changes in adipose tissues metabolism may be the result of the differences in gene expression pattern in female and male offspring in response to MO. DEG analysis revealed that very few genes (60) were differently expressed between SAT and VAT in M-moHF, as opposed to females that showed about 1,600 DEG between SAT and VAT. These differences in gene expression could indicate that males would not differentiate the two white adipose depots which, in turn, will promote VAT growth and local inflammation, as opposed to females. A number of studies demonstrated that MO promotes inflammation and insulin resistance in F1 offspring, especially in males (Chang et al., 2019; Litzenburger et al., 2020). In the current study, we showed that males undergo an important remodeling in WAT metabolism, especially in SAT, in line with others (Chang et al., 2019), as opposed to females that showed important transcriptional modifications in BAT metabolism. These results would support the damaging effect of MO in males WAT, on the long term. Reduced ω6:ω3 ratio in WAT has been associated with improved lipid metabolism, reduced inflammation and oxidative stress (Yang et al., 2016). In our study, females showed reduced ω6:ω3 ratio as opposed to males that induced it in response to MO. Most interestingly, we found a large number of biological pathways that were oppositely regulated between sexes and by MO with key regulated genes. Female and male offspring had sex-dependent transcriptional adaptation to MO that may oppositely drive obesity and its associated metabolic complications later in life between sexes. In line with this statement, MO has been shown to modulate two major genes involved in adipogenesis (*Sfrp4*) (Zhang et al., 2020) and in TG synthesis (*Dgat2*) (Chitraju et al., 2019) in a sex-dependent manner in VAT. Female offspring born from obese mothers promoted *Fabp4* expression in SAT, which function as a positive factor in fatty acid signaling (Hertzel and Bernlohr, 1998), and induced BAT activity by repressing *Kng2* (blunt thermogenesis) expression (Peyrou et al., 2020) and inducing *Elovl3* and *Mgll* (FA elongation and degradation) expression levels as opposed to males. On the other hand, female but not male offspring born from obese mothers showed reduced Homa index as compared to those born from lean mothers, associated to reduced level of C16:1 FA, a circulating lipokine that modulate insulin sensitivity in the liver (Cao et al., 2008). Although the precise mechanism remains to be established, our results suggest that MO differently reprograms adipose tissues in female and male offspring. One can speculate that MO impacts female offspring metabolism differently than males by reprogramming adipose tissue in females towards more energy dissipation (Increased *Ucp1* and decreased *Kng2*) and lipid oxidation (increased *Noct, Acad11* and *Ephx2*) as opposed to males, and by reprogramming adipose tissue in males towards insulin sensitivity (reduced *Fapb4/5*) and SAT inflammation (induced *Ccl5/12/19* and *Cxcl12/15*).

In conclusion, female and male offspring on obesogenic diet show different metabolic outcomes with MO due to different remodeling of the transcriptome in VAT, SAT and BAT. These sex-dependent observations in adipose tissue biological pathways may be a key contributor to the sexual dimorphism we observed in the offspring’s lipidome in response to MO. These novel findings may help to better prevent metabolic alterations in offspring and strongly support the concept that adipose depots have different metabolic functions due to different transcriptional regulation. In addition, we present further evidence that male and female offspring metabolic adaptation to MO occurs in a sex-dependent and adipose depot-dependent manner, which may set the basis for targeted medicine.

## EXPERIMENTAL MODEL AND METHOD

### Mice and diet

All animal procedures were approved by the local Ethical Committee of the Swedish National Board of Animal Experiments. Four-week-old virgin dams and sire C57Bl6/J were ordered and recovered for one week before F0 dams were randomized to control diet (CD; D12450H, Research Diets, NJ, USA; 10% kcal fat from soybean oil and lard; n=6, F0-CD) or to high fat diet (HFD; D12451, Research Diets, NJ, USA; 45% kcal fat from soybean oil and lard; n=6, F0-HFD) for six weeks before mating. Sires remained on CD until mating. After six weeks of their respective diet two F0 dams were mated with one F0 sire. During this short mating period (up to five days) sires were on the same HFD as dams in the group (experimental unit). We assumed that the sires spermatozoa were unlikely affected by the HFD while mating as sperm maturation time is approximately 35 days (Oakberg, 1956). After mating, F0 sir and pregnant dams were separated. F0 dams were continuously exposed to their respective diets throughout pregnancy and until weaning. The F1 offspring were weaned at postnatal day 21 (3-week). Afterwards, F1 male and female offspring were separated, three to five animals were randomly housed per cage and fed with the HFD until the end of the study (Fig. 1A). The group of offspring born from HFD fed dams were named moHF and the group of offspring born from CD fed dams were named moC. All mice were housed in a 23°C temperature-controlled 12 h light/dark room, with free access to water and food unless specified. Body weight (BW) was recorded at midterm (MID, 15 weeks) and at endterm (END, 26 weeks) timepoints of the study in all groups. The average food intake (per cage) in offspring was recorded twice a week for two weeks at around 4-month of age.

### In vivo magnetic resonance imaging (MRI)

Animals were anesthetized using isoflurane (4% for sleep induction and ~2% for sleep maintenance) in a 3:7 mixture of oxygen and air, before being positioned prone in the MR-compatible animal holder. Respiration was monitored during scanning (SA-instruments, Stony Brook, NY, USA). Core body temperature was maintained at 37°C during scanning using a warm air system (SA-instruments, Stony Brook, NY, USA).

The magnetic resonance imaging (MRI) experiments (n=4-7 per group) were conducted on the same mouse at week 14 and week 25 using a 9.4 T horizontal bore magnet (Varian Yarnton UK) equipped with a 40 mm millipede coil, as previously described (Korach-Andre, 2020). Fiji software (http://fiji.sc) was used to compute the volume of total fat (TF), visceral adipose tissue (VAT), subcutaneous adipose tissue (SAT) and brown adipose tissue (BAT).

### In vivo localized proton magnetic resonance spectra (^1^H-MRS)

As for the MRI scanning, animals were anesthetized before being positioned prone in the MR-compatible animal holder. Respiration was monitored and body temperature maintained at 37°C during scanning. In addition, heart beats were recorded using an electrocardiogram system as previously described (Korach-Andre, 2020). Localized ^1^H-MRS from visceral and subcutaneous fat depots were acquired from 2 x 1.5 x 1.5 mm^3^ voxels positioned in the upper gonadal abdominal fat (as representative of visceral fat) and in the inguinal abdominal fat (as representative of the subcutaneous fat) (Fig.2A). Point Resolved Spectroscopy (PRESS) was used as primary pule sequence (Strobel et al., 2008) with the following parameters: time to echo 15 ms, sweep width 8013 Hz, number of excitations 16, refocusing pulses 1.6 ms mao pulses with a band width nominal bandwidth of 2936 Hz as described (Korach-Andre, 2020).

All spectroscopy data were processed using the LCModel analysis software (http://s-provencher.com/pub/LCModel/manual/manual.pdf). “Lipid 6” for adipose spectrum were used as a base with all signals occurring in the spectral range of 0 to 7 ppm (water resonance at 4.7ppm) simulated in LCModel. All concentrations were derived from the area of the resonance peaks of the individual metabolites. Only the fitting results with an estimated standard deviation of less than 20% were further analyzed. ^1^H-MRS spectra revealed nine lipid signals (peaks) in the mouse adipose, based on published data (Mosconi et al., 2014; Ye et al., 2012). As for the MRI, ^1^H-MRS experiments were repeated twice on the same animal at MID and END.

### Biochemical analysis of plasma

Glucose and insulin levels were measured after 6 h fasting from 7 a.m. to 1 p.m. Glucose level was measured instantly via tail-nick one-touch glucometer. Extra blood was collected using a capillary tube for insulin measurements in plasma using a commercial rat/mouse Insulin Elisa kit (EZRMI-13K) according to the manufacturer instructions. Matsuda index (whole body insulin sensitivity index) and direct measurement of hepatic insulin resistance (HOMA index) were calculated as described (Matsuda and DeFronzo, 1999; Pacini et al., 2013). Briefly, homeostatic model assessment (HOMA) index was calculated as follows = (I_0_ x G_0_)/22.5. Matsuda index was calculated as = 100/(✓[G_0_ x I_0_ x G_mean_ x I_mean_]), the suffix *mean* indicates the average value of glucose and insulin concentration measured during the whole length of the glucose test. Evaluation of β-cell function was calculated by dividing the area under the curve (AUC) of insulin and glucose levels during the glucose test (AUCins:AUCglc).

### Fatty acid analysis using gas chromatography with a flame ionization detector (GC-FID)

Total lipid extracts were obtained using a modified Bligh and Dyer method (Folch et al., 1957) and after transmethylation, the fatty acids were analyzed by gas chromatography with a flame ionization detector (GC-FID) using a modification of the method of Aued-Pimentel et al. (Aued-Pimentel et al., 2004). Fatty acid methyl esters (FAMEs) were dissolved in 30 μL of *n*-hexane and 2.0 μL were injected in GC-FID (PerkinElmer Clarus 400 gas chromatograph (Waltham, MA). The gas chromatograph injection port was programmed at 215°C and the detector at 250°C. The initial temperature was 75°C and the oven temperature was programmed in 3 ramps (a 15°C/min increase to 163°C for 2 min, a 2°C/min increase to 175°C for 2 min, and a 10°C/min increase to 250°C form 5 min), performed for 28.3 min in total. Hydrogen was the carrier gas (flow rate, 1.7 ml/min). A DB-FFAP column (30m long, 0.32 mm internal diameter, and 0.25 μm film thickness (J & W Scientific, Folsom, CA, USA)) was used. Peaks corresponding to each FA were identified based on retention time in comparison with a Supelco 37 Component FAME standard mixture (Sigma-Aldrich, USA), integrated and the percentage of each FA was related to the sum area of all FAs identified. The total ω-3 content was calculated as the summed total of ω-3 PUFA of C18:3ω-3 and C20:3ω-3. Total ω-6 content was calculated as the summed total of C18:2ω-6 and C20:3ω-6 contents.

### LC-MS analysis of triglycerides

Total lipid extracts were separated using a high-performance liquid chromatography (HPLC) system (Ultimate 3000 Dionex, Thermo Fisher Scientific, Bremen, Germany) with an autosampler coupled online to a Q-Exactive hybrid quadrupole Orbitrap mass spectrometer (Thermo Fisher Scientific, Bremen, Germany), adapted from (Anjos et al., 2019; Colombo et al., 2018) LC-MS analysis was carried out using an Accucore™ C30 column (150 × 2.1 mm) that was equipped with 2.6 μm diameter fused-core particles (Thermo Fisher Scientific, Germering, Germany). The solvent system consisted of two mobile phases: mobile phase A (water/ACN 50/50 (v/v) with 0.1% formic acid and 5 mM ammonium formate) and mobile phase B (isopropanol/ACN/water 85/10/5 (v/v) with 0.1% formic acid and 5 mM ammonium formate). Initially, 50% of mobile phase B was held isocratically for 2 min, followed by a linear increase to 86% of B within 18 min. An increase to 95% B occurred in 1 min, which was held up for 14 min, returning to the initial conditions (50% B) in 2 min, followed by a re-equilibration period of 8 min prior to the next injection. An aliquot of 20 μg of each lipid extract were dissolved in 80 μL of MeOH. Two μL of each dilution were introduced into the Accucore™ C30 column (150 × 2.1 mm) that was equipped with 2.6 μm diameter fused-core particles (Thermo Fisher Scientific, Germering, Germany) with a flow rate of 300 μL min^-1^. The temperature of the column oven was maintained at 40°C. The mass spectrometer with Orbitrap technology operated in positive (electrospray voltage 3.0 kV) ion mode with a capillary temperature of 350°C, a sheath gas flow of 45 arbitrary units (a.u), an auxiliary gas flows of 15 a.u., a high resolution of 70 000, a maximum injection time of 100 ms and AGC target 1e6. In MS-MS experiments, cycles consisted of one full scan mass spectrum and ten data-dependent MS-MS scans (resolution of 17 500, a maximum injection time of 100 ms an AGC target of 1e5, and with an isolation window of 1 *m/z*). Cycles were repeated continuously throughout the experiments with the dynamic exclusion of 60 s and an intensity threshold of 5e4. Normalized collisional energy ranged between 20, 23, and 25 eV. C30 RP-LC-MS spectra from molecular species of triglyceride were analyzed in positive ion mode and TG were identified as [M+NH_4_Γ ions. Data acquisition was carried out using the Xcalibur data system (V3.3, Thermo Fisher Scientific, USA). The mass spectra were processed and integrated through the MZmine software (v2.32) (Pluskal et al., 2010). This software allows for filtering and smoothing, peak detection, alignment and integration, and assignment against an in-house database, which contains information on the exact mass and retention time for each TG molecular species. During the processing of the data by MZmine, only the peaks with raw intensity higher than 1e4 and within 5 ppm deviation from the lipid exact mass were considered. The identification of each TG species was validated by analysis of the MS/MS spectra. The MS/MS spectra of [M+NH_4_Γ ions of TGs allowed the assignment of the fatty acyl substituents on the glycerol backbone (Hsu and Turk, 2010).

### RNA isolation, purity and integrity determination

Total RNA was extracted from VAT, SAT and BAT using QIAGEN miRNeasy Mini Kit (217004, Qiazol) and RNase-Free DNase Set (79254) for removing possible DNA traces. RNA concentration was measured by nanodrop and diluted to 2.17 ng/μl. RNA quality was assessed for 12 out of 39 samples randomly by a 2100 Bioanalyzer using a nano chip (Agilent Biotechnologies), and RIN numbers were between 8.7-9.9. cDNA libraries were prepared for the bulk-RNA sequencing in 384-well plate with triplicates for each RNA sample according to the previously described Smart-Seq2 protocol (Picelli et al., 2014). In brief, mRNA was transcribed into cDNA using oligo(dT) primer and SuperScript II reverse transcriptase (ThermoFisher Scientific). Second strand cDNA was synthetized using a template switching oligo, followed by PCR amplification for 15 cycles. Purified cDNA was quality controlled on a 2100 Bioanalyzer with a DNA High Sensitivity chip (Agilent Biotechnologies), fragmented and tagged (tagmented) using Tn5 transposase, and each single sample was uniquely indexed using the Illumina Nextera XT index kits (Set A-D). Subsequently, the libraries were pooled into one lane for sequencing at a HiSeq3000 sequencer (Illumina), using sequencing format of dual indexing and single 50 base-pair reads.

### Bulk RNA-seq mapping

All raw sequence reads available in FastQ format was mapped to the mouse genome (mm10) using Tophat2 with Bowtie2 option (Kim et al., 2013; Langmead and Salzberg, 2012), where adaptor sequences were removed using trim galore before read mapping. BAM files containing the alignment results were sorted according to the mapping position. Raw read counts for each gene were calculated using featureCounts from Subread package (Liao et al., 2014).

### Bulk RNA-seq differential gene expression analysis

DEseq2 was used to perform the analysis of differential gene expression, where genes with raw counts as input (Love et al., 2014). The differentially expressed genes were identified by adjust p value for multiple testing using Benjamini-Hochberg correction with False Discovery Rate (FDR) values less than 0.1.

### Pathway analysis

The pathway analysis, also called Gene Set Enrichment Analysis (GSEA) (Mootha et al., 2003), was performed using the KEGG pathways dataset. First, genes were ranked descending according to the Log2 Fold Change (Log2FC) of expression. For each query pathway, if gene *i* is a member of the pathway, it is defined as

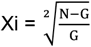

If gene *i* is not a member of the pathway, it is defined as

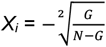

where *N* indicates the total number of genes and *G* indicates the number of genes in the query pathway. Next, a max running sum across all *N* genes Maximum Estimate Score (MES) is calculated as

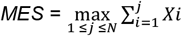

The permutation test was performed with 1000 times to judge the significance of MES values. The query pathway with a nominal p-value less than 0.05 and FDR values less than 0.1 would be considered to be significantly enriched. The positive MES value indicates up-enrichment (up-regulation) whereas a negative MES value indicates down-enrichment (down-regulation) of a pathway.

### Statistical analyses

Data are expressed as means ± standard error of the mean (Weigt et al.). Differences between offspring sex and mother diet groups (F-moC, M-moC, F-moHF and M-moHF) were determined using two-way ANOVA with diet (D) and sex (S) as independent variables, followed by Tukey’s multiple comparison post hoc test when significant (p<0.05). Differences between two groups (sexes, F *versus* M; maternal diet moC *versus* moHF) were determined by t-test corrected for multiple comparisons using the Holm-Sidak method, with alpha=5.000%. ^*^, p<0.05 M *versus* F and ^#^, p<0.05 moHF *versus* moC within the same sex were considered significant. ^**^ or ^##^, p<0.01; ^***^ or ^###^, p<0.001.

## Supporting information

Metabolic pathways in VAT, SAT and BAT

TG species extracted from BAT

Fatty acids contained in TG extracted from BAT

Significant DEG between sexes in VAT, SAT and BAT of female and male offspring born from lean and obese mothers

Significant DEG in response to maternal obesity in VAT, SAT and BAT of female and male offspring

## Author Contributions

M.K.A. conceptualized and designed the study. C.S., M.G.G and M.K.A. performed animal experiments; C.S. and M.K.A. collected and analyzed data; L.H., D.C., T.M. and R.M.D. performed and analyzed lipidomic data, wrote the method for lipidomic; B.B., S.G. and J.L. performed RNA sequencing experiments; X.L. performed the bioinformatics; C.S. and M.K.A. designed the figures and drafted the manuscript, which was substantially edited and approved by all authors. M.K.A is the guarantor of this work and, as such, had full access to all the data in the study and take responsibility for the integrity of the data and the accuracy of the data analysis. All authors approved the final version of the manuscript.

## Acknowledgements

The MRI and MRS experiments were performed at the Department of Comparative Medicine/Karolinska Experimental Research and Imaging Centre at Karolinska University Hospital, Solna, Sweden. We thank Peter Damberg and Sahar Nikkhou Aski for excellent assistance to develop the sequence for proton-magnetic resonance spectroscopy in the adipose tissue. We thank Vasiliki Karagianni for excellent technical assistance for animal work. This work and M.K.A. were supported by the Novo Nordisk Foundation (NNF14OC0010705), by the Lisa and Johan Grönbergs Foundation (2019-00173) and by AstraZeneca (ICMC). L.A.H. is supported by grants from FCT-Fundação para a Ciência e a Tecnologia (UID/BIM/04501/2020), CCDRC (CENTRO-01-0145-FEDER-000003) and CCDRC (CENTRO-01-0246-FEDER-000018), M.R.D. is supported by CESAM (UIDP/50017/2020+UIDB/50017/2020), QOPNA (FCT UID/QUI/00062/2019) and LAQV/REQUIMTE (UIDB/50006/2020).

---

**Supplementary Figure S1. Metabolic pathways in VAT, SAT and BAT**.

(A) Clustered heatmap of all KEGG pathway enrichment analysis presenting the expression level between female and males (sex) and between moC and moHF (diet) in VAT (n=3-6).

(B) Clustered heatmap of all KEGG pathway enrichment analysis presenting the expression level between female and males (sex) and between moC and moHF (diet) in SAT (n=3-6).

(C) Clustered heatmap of all KEGG pathway enrichment analysis presenting the expression level between female (F) and males (M) (sex) and between moC and moHF (diet) in BAT (n=3-6).

**Supplementary Figure S2. TG species extracted from BAT**.

(A) Plot of the low abundant TG species in BAT detected by LC-MS (n=4).

(B) Plot of the moderate abundant TG species in BAT detected by LC-MS (n=4).

(C) Plot of the high abundant TG species in BAT detected by LC-MS (n=4).

**Supplementary Figure S3. Fatty acids contained in TG extracted from BAT**.

(A) Plot of the FA species in BAT detected by GC-MS (n=4).

(B) The ω-3 and ω-6 FA pathways and the ratio of ω-6 to ω-3 FA pathways.

(C) Pie charts of the FA saturation profile.

**Supplementary Table S1. Significant DEG between sexes in VAT, SAT and BAT of female and male offspring born from lean and obese mothers**.

**Supplementary Table S2. Significant DEG in response to maternal obesity in VAT, SAT and BAT of female and male offspring**.

