## Supplementary material for "Maternal obesity programs white and brown adipose tissue transcriptome and lipidome in offspring in a sex-dependent manner": Metabolic pathways in VAT, SAT and BAT

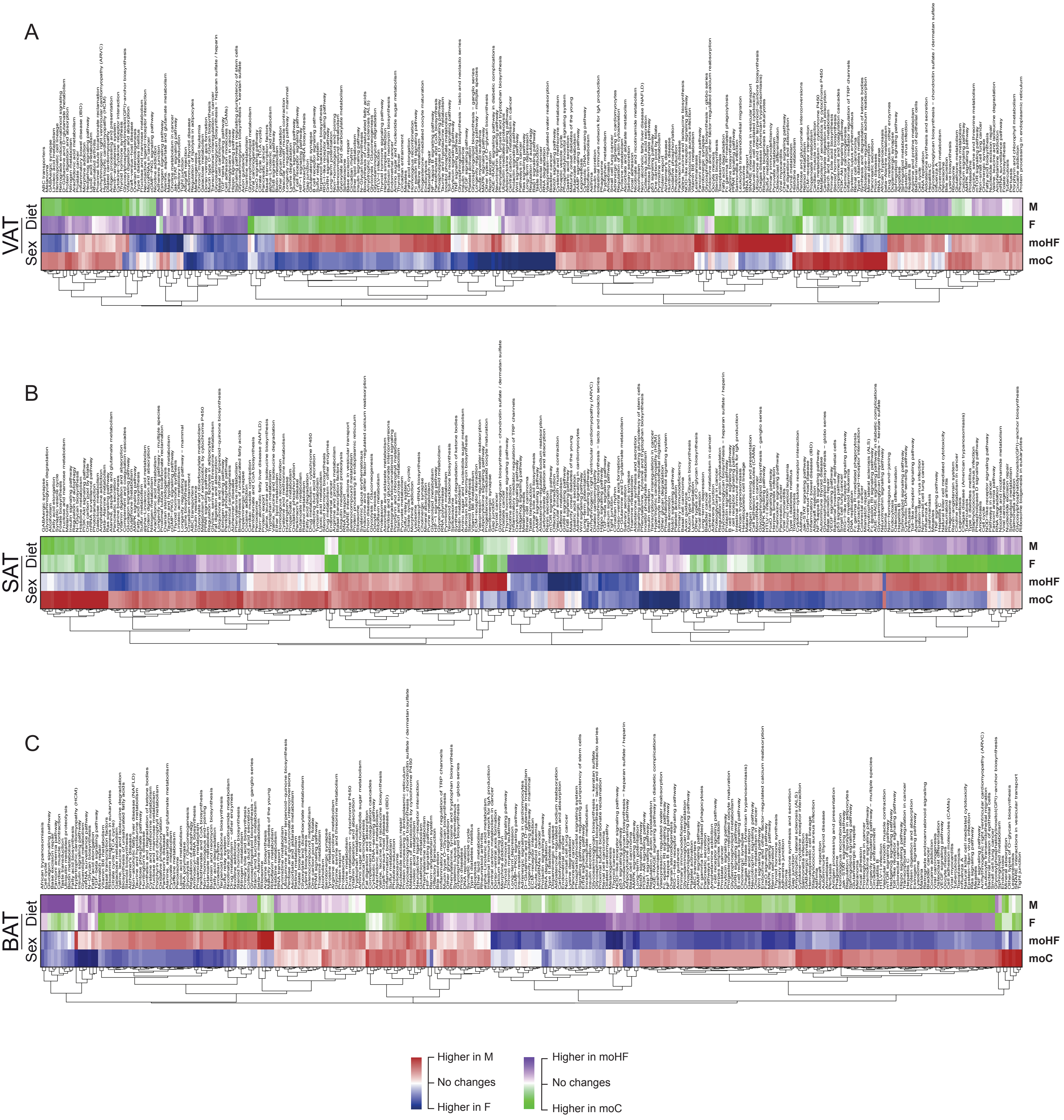


**Supplementary Figure S1. Metabolic pathways in VAT, SAT and BAT.** (A) Clustered heatmap of all KEGG pathway enrichment analysis presenting the expression level between female and males (sex) and between moC and moHF (diet) in VAT (n=3-6). (B) Clustered heatmap of all KEGG pathway enrichment analysis presenting the expression level between female and males (sex) and between moC and moHF (diet) in SAT (n=3-6).(C) Clustered heatmap of all KEGG pathway enrichment analysis presenting the expression level between female and males (sex) and between moC and moHF (diet) in BAT (n=3-6).
