## Supplementary material for "Maternal obesity programs white and brown adipose tissue transcriptome and lipidome in offspring in a sex-dependent manner": TG species extracted from BAT

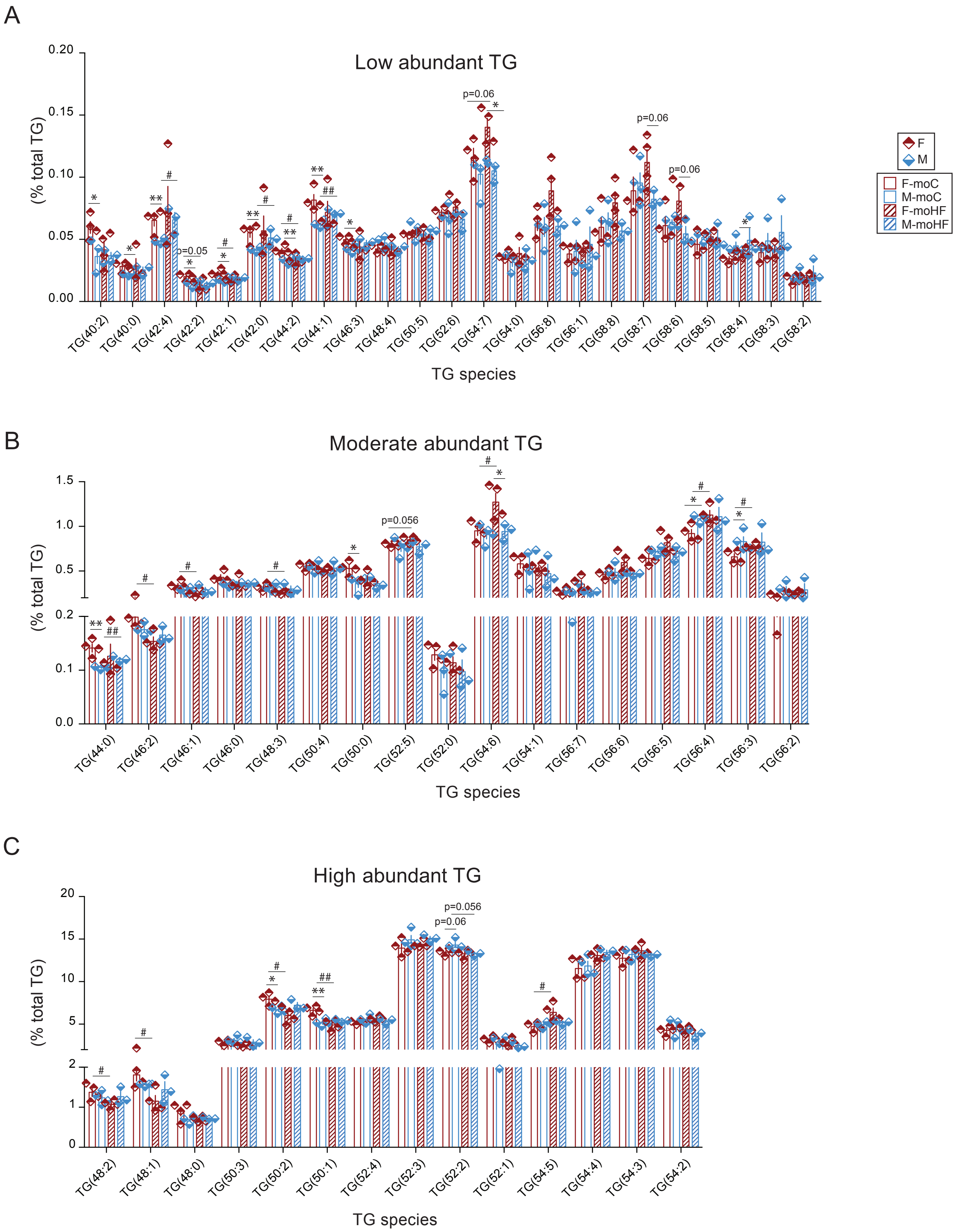


**Supplementary Figure S2. TG species extracted from BAT.**
(A) Plot of the low abundant TG species in BAT detected by LC-MS (n=4).
(B) Plot of the moderate abundant TG species in BAT detected by LC-MS (n=4).
(C) Plot of the high abundant TG species in BAT detected by LC-MS (n=4).
