## Supplementary material for "Maternal obesity programs white and brown adipose tissue transcriptome and lipidome in offspring in a sex-dependent manner": Fatty acids contained in TG extracted from BAT

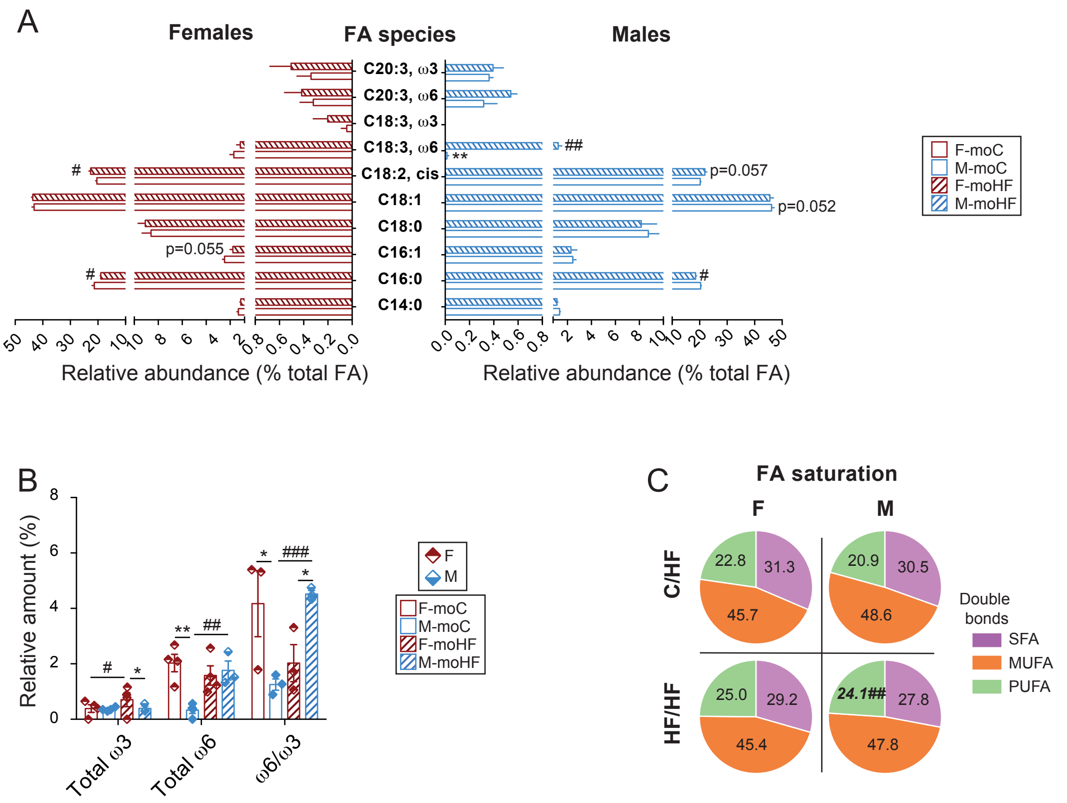


**Supplementary Figure S3. Fatty acids contained in TG extracted from BAT.**
(A) Plot of the FA species detected by GC-MS (n=4) in BAT
(B) The ω−3 and ω−6 FA pathways and the ratio of ω−6 to ω−3
FA pathways. (C) Pie charts of the FA saturation profile.
