## Supplementary material for "Maternal obesity programs white and brown adipose tissue transcriptome and lipidome in offspring in a sex-dependent manner": Significant DEG between sexes in VAT, SAT and BAT of female and male offspring born from lean and obese mothers

Suppl.Table S1: Significant DEG between sexes in VAT, SAT and BAT of female and male offspring born from lean and obese mothers.

| | **Males *versus* Females** | | |
|---|---|---|---|
| | VAT | SAT | BAT |
| **moC** | **Inflammation**: Tnfrsf11a; Rock2; Gnb4; Blc2a1b; Rap1b; Adcy3; Ccr1; Dock2; Ccl9: Adcy7; Itgb2; Ifngr1; Cxcl13; Gnb5<br>**Insulin/glucose**: Ppp1cc; Pde3b; Prkar2b; Acacb; Hk3; Aldh1a3; Flot1; Eno1; Kcnn4; Adcy3; Adcy7<br>**Oxidative phosphorylation**: Tcirg1; Atp6v0d2; Atp6v0e<br>**Lipid metabolism**: Adcy3; Adcy7; Plin1; Pde3b; Ptger3; Elovl3; Elovl6; Fads1; Angptl4; Ilk; Fabp5; Pparg; Abca1; Abca8a; Acacb | **Inflammation**: Prkcq; Malt1; Ly96; Ltb; Tnfrsf13c; Gadd45b; Cxcl9; Cxcl12; Cxcl15; Ccl11; Ccl21a; Cxcr3; Cxcr5; Cxcr6; Ccr9; Stat3; Prkaca; Hck; Akt2; Itk; Gng2; Gng5; Gng11; Gngt2; Dock2; Prkcd; Grk6; Pak1; Lck; Ptpn6; Vav1; Rac2; Rac1; Itgal; Lat; Plcg2; Ifngr1; Ifnar2<br>**Insulin/glucose**: Calm1; Calm3; Pde3b; Prkar2b; Flot2; Sorbs1; Eif4ebp1; Rheb; Mknk2; Atp1a2; Atp1b1; Prkaca; Rps6ka1; Ptprf; Stat3; Gys1; Gys2; Acacb; Slc27a1; Cd36; Prkcq; Prkcd; Nr1h3; Akt2; Ogt; Slc2a1; Adh1; Pgm1; Dld; Tpi1; Dlat; Pfkl; Eno3; Adh5; Pdhb; Aldoc; Gapdh; Ldha; Fbp2<br>**Oxidative phosphorylation**: mt-Nd2; Ndufs3; mt-Nd4; mt-Nd1; mt-Nd5; mt-Nd3; Ndufs1; mt-Nd6; Ndufs4; Ndufs5; Ndufs6; Ndufs7; Ndufs8; Ndufv1; Ndufv2; Ndufv3; Ndufa1; Ndufa2; Ndufa3; Ndufa4; Ndufa5; Ndufa6; Ndufa7; Ndufa8; Ndufa9; Ndufa10; Ndufab1; Ndufa11; Ndufa12; Ndufa13; Ndufb2; Ndufb3; Ndufb4; Ndufb5; Ndufb6; Ndufb7; Ndufb8; Ndufb9; Ndufb10; Ndufb11; Ndufb11; Ndufc1; Ndufc2; Uqcrfs1; mt-Cytb; Cyc1; Uqcrc1; Uqcrc2; Uqcrh; Uqcrb; Uqcrq; Uqcr10; Uqcr11; mt-Co1; mt-Co2; Cox4i1; Cox5a; Cox5b; Cox6a1; Cox6b1; Cox6c; Cox7a2; Cox7b; Cox7c; Cox8a; Cox8b; Cox17; Atp5a1; Atp5b; Atp5c1; Atp5d; Atp5e; Atp5o; mt-Atp6; Atp5pb; Atp5g2; Atp5g3; Atp5h; Atp5k; Atp5j2; Atp5l; Atp5j; mt-Atp8; Atp6v1b1; Atp6v1e1; Atp6v1f; Atp6v0b; Atp6v0e; Atp6v0e2; Ppa1; Ppa2; Mdh1; Idh3g; Sdhd; Pcx; Sucla2; Idh3a; Idh3b; Sdhc; Mdh2; Suclg1; Dld; Sdhb; Fh1; Dlat; Sdha; Pdhb; Dlst<br>**Lipid metabolism**: Mgll; Abhd5; Prkaca; Pnpla2; Pde3b; Plaat3; Adora1; Elovl7; Adh1; Eci1; Hadhb; Acadvl; Echs1; Hadh; Eci2; Acaa2; Acads; Adh5; Acadsb; Hsd17b12; Elovl6; Fads1; Hacd2; Hadha; Acot1; Hacd1; Angptl4; Acadl; Scp2; Fabp5; Fabp4; Acox1; Ehhadh; Plin2; Dbi; Aqp7; Slc27a1; Me1; Cpt2; Sorbs1; Acadm; Lpl; Nr1h3; Plin4; Pltp; Plin5; Abcd3; Abcb6; Abcg2; Akt2; Cd36; Acsl1; Prkcq; Lepr; Stat3; Rxrg; Adipoq; Acacb; Slc2a1 | **Inflammation**: Tyrobp; Fcer1g; Cxcl14; Ccl11<br>**Insulin/glucose**: Hk2; Acss2; Pdha1; Irs1; Ppp1cb; Ppp1r3b; Acaca; Prkag2; Gnas; Cd36<br>**Oxidative phosphorylation**: Ndufa3; Ndufc1; Cox7a1; Cox7c; Cox8b; Atp5k; Ppa1; Cs; Pdha1<br>**Lipid metabolism**: Adrb3; Mgll; Gnas; Acaca; Hacd2; Ehhadh; Lpl; Angptl4; Acox1; Plin2; Ubc; Abcd2; Irs1; Cd36; Acsl5; Prkag2 |
| **moHF** | **Inflammation**: Tnfrsf1a; Itgb2; Ifngr1; Mmp9<br>**Insulin/glucose**: Ptprf; Atp1b1; Tnfrsf1a<br>**Oxidative phosphorylation**: Atp6v0a1; Atp6v0a4<br>**Lipid metabolism**: Elovl1; Hacd2; Acot2; Tnfrsf1a | **Inflammation**: Cxcl15; Ifngr1<br>**Insulin/glucose**: Ptprf<br>**Oxidative phosphorylation**: Atp6v1b1<br>**Lipid metabolism:** Fads1 | **Inflammation**: Cflar<br>**Insulin/glucose**: Acss1; Aldob; Aldoa; Pdha1; Gys1; Foxo1<br>**Oxidative phosphorylation**: Ogdh; Pdha1<br>**Lipid metabolism**: Plin2; Fabp1; Apoa2; Hmgcs2 |

DEG analysis showing the significantly changed genes that belong to the selected KEGG pathways (inflammation, Insulin/glucose, oxidative phosphorylation and lipid metabolism) and that were up-(fold2change>0) and down-(fold2change<0) regulated by sex in VAT, SAT and BAT of offspring born from control mothers (moC) and obese mothers (moHF) (n=3 to 6) using FDR<0.1 and p-value< 0.05 as a cut-off.
