## Supplementary material for "Maternal obesity programs white and brown adipose tissue transcriptome and lipidome in offspring in a sex-dependent manner": Significant DEG in response to maternal obesity in VAT, SAT and BAT of female and male offspring

Suppl.Table S2: Significant DEG by maternal obesity in VAT, SAT and BAT of female and male offspring.

| | **moHF *versus* moC** | | |
|---|---|---|---|
| | VAT | SAT | BAT |
| FEMALES | **Inflammation**: Ccl7; Ccl11; Rock2; Gnb5<br>**Insulin/glucose**: Atp1a2; Adcy3; Gys2; Slc27a1; Insr; Pck1; Aldoa; Ldha; Tpi1<br>**Oxidative phosphorylation**: Pck1<br>**Lipid metabolism**: Adhd5; Insr; Ucp1; Slc27a1; Fabp1; Apoa2; Abca1; Pck1 | **Inflammation**: -<br>**Insulin/glucose**: Pde3b; Pck1<br>**Oxidative phosphorylation**: Pck1<br>**Lipid metabolism**: Pck1; Fabp4; Pde3b | **Inflammation**: Cebpb; Tyrobp; Cxcl12; Ccl9; Adcy3<br>**Insulin/glucose**: Dlat; Acss2; Eno1; Aldoa; Pdhb; Aldh9a1; Gapdh; Ldha; Adh5; Ppp1cb; Pygl; Acaca; Prkab1; Atp1a2; Adcy3<br>**Oxidative phosphorylation**: Ndufa6; Sdhd; Cox7b; Ppa1; Mdh1; Pdhb; Dlat<br>**Lipid metabolism**: Me; Lpl; Ucp1; Acox1; Dbi; Pltp; Mgll; Adcy3; Adrb3; Acox2; Acaca; Aldh9a1; Acat2; Adh5; Elovl3; Acot2; Prkab1 |
| MALES | **Inflammation**: Tnfrsf11a<br>**Insulin/glucose**: Aldob; Atp1b1; Slc27a5; Slc27a2; Tnfrsf1a<br>**Oxidative phosphorylation**: Atp6v0e<br>**Lipid metabolism**: Plin1; Acsl3; Elovl2; Acot2; Apoa1; Apoa2; Slc27a5; Apoa5; Apoc3; Fabp1; Slc27a2; Tnfrsf1a | **Inflammation**: Prkcq; Btk; Card11; Birc2; Birc3; Cd40; Lta; Ltb; Tnfrsf13c; Bcl2a1b; Vcam1; Gadd45b; Ccl21a; Cxcr6; Ccl6; Elmo1; Ccl12; Cxcr5; Pf4; Ccl5; Dock2; Ccl19; Cxcl16; Fgr; Ccl24; Ptpn6; Itgal; Klrd1; Lck; Cd247; Lat<br>**nsulin/glucose**: Gys2; Nos3; Prkcq; Ogt; Eno3; Pkm<br>**Oxidative phosphorylation**: mt-Nd2; Ndufs4; Ndufs7; Ndufs8; Ndufv2; Ndufa2; Ndufa4; Ndufa6; Ndufab1; Ndufa11; Ndufa12; Ndufa13; Ndufb2; Ndufb3; Ndufb5; Ndufb6; Ndufb7; Ndufb8; Ndufb9; Ndufb11; Ndufc2; Uqcrb; Uqcrq; Uqcr10; Uqcr11; Cox4i1; Cox5a; Cox5b; Cox6a1; Cox6b1; Cox7c; Cox8a; Cox8b; Atp5d; Atp5o; Atp5g3; Atp5h; Atp5j2; Atp5l; Atp5j; Atp6v0e<br>**Lipid metabolism**: Hmgcs2; Fabp5; Fabp4; Dbi; Abcg2; Prkcq; Lepr | **Inflammation**: -<br>**Insulin/glucose**: Hk2; Pdha1<br>**Oxidative phosphorylation**: Ndufa3; Cox7a1; Cox8b; Atp5k; Idh3a; Pdha1<br>**Lipid metabolism**: Acsl5 |

DEG analysis showing the significantly changed genes that belong to the selected KEGG pathways (inflammation, Insulin/glucose, oxidative phosphorylation and lipid metabolism) and that were up-(fold2change>0) and down-(fold2change<0) regulated by maternal diet in VAT, SAT and BAT of female (F) and male (M) (n=3 to 6) offspring using FDR<0.1 and p-value< 0.05 as a cut-off.
